# Bulk segregant analysis reveals genomic regions associated with imidacloprid resistance in the Colorado potato beetle

**DOI:** 10.1101/2025.01.20.633857

**Authors:** Alitha Edison, Nijat Narimanov, Pablo Duchen, Shuqing Xu

**Affiliations:** Institute for Evolution and Biodiversity, University of Münster, Münster, Germany; Institute of Organismic and Molecular Evolution (iomE), University of Mainz, Mainz, Germany

**Keywords:** insecticide resistance, QTL mapping, neonicotinoid, pesticide, *Leptinotarsa decemlineata*, genetic basis of resistance RNA interference

## Abstract

Given the increased accessibility of genomic techniques, the speed of evolution of resistance, and the large number of genes involved in resistance, investigations into the genetic basis of resistance in more species are pertinent. Despite being an important agricultural pest, only a limited number of genetic mapping studies based on crossed populations have been performed to identify genes involved in resistance in the Colorado potato beetle (*Leptinotarsa decemlineata*, CPB). Here, we performed bulk segregant analysis on a mapping population generated by creating advanced intercross lines from five European strains of CPB. We identified eight peaks across chromosomes 1,8,10, and 16 involved in resistance against the neonicotinoid, imidacloprid. We identified 337 genes within these peaks and shortlisted three candidates based on gene expression and functional annotation. Among the three candidates thus identified, we found that an ABC transporter and a galactosyl transferase are expressed in relatively higher amounts in a relatively more susceptible strain than in a resistant strain. We attempted to validate the role of these two genes in insecticide resistance by knocking them down in a resistant strain using RNA interference (RNAi) and performing toxicity experiments. Surprisingly, our results showed that the activation of RNAi machinery reduced imidacloprid resistance and the effect is not specific to the tested candidate genes, which raised a concern on the suitability of using RNAi for validating insecticide resistance mechanisms in CPB.

## 1. Introduction

Insecticide resistance is a classic example of natural selection and a global economic problem threatening sustainable food production (Hawkins et al., 2019). Understanding the genetic basis of resistance is crucial for both academic research and applied pest management. This is especially relevant in the current environment where research on genetics- and genomics-based pest management is rapidly advancing (Li et al., 2022; Palli, 2014; Y. Yan et al., 2023).

Several target site-associated genes and metabolic genes have been shown to be involved in insecticide resistance. Mutations in conventional target sites, including neurotransmitter receptors, and ligand- and voltage-gated ion channels, have been shown to increase resistance (ffrench-Constant et al., 1993; Mutero et al., 1994; Williamson et al., 1996). Among genes that are involved in important physiological functions like xenobiotic detoxification, transport and excretion in insects, the most ubiquitous resistance-associated candidates are cytochrome P450 monooxygenases (CYPs), esterases (ESTs), ATP-binding cassette transporters (ABCs) and glutathione S-transferases (GSTs) (Amezian et al., 2024; Enayati et al., 2005; Hemingway, 2000; Liu et al., 2015; Nauen et al., 2022; Pavlidi et al., 2018). In addition to mutations in target sites and metabolic genes, mutations causing upregulation and downregulation of genes by both cis- or trans-regulatory processes have also been shown to be involved with resistance. Novel mechanisms (L. P. Chen et al., 2023; Pu and Chung, 2024) are being discovered, possibly faster than ever before, owing to technical advances in genetic mapping and reduced costs of sequencing.

Colorado potato beetles (CPB, *Leptinotarsa decemlineata*) are highly destructive leaf-eating pests of potato (*Solanum tuberosum*). CPB is currently resistant against more than 50 different types of insecticides like DDT, imidacloprid, thiamethoxam and spinosad (Mota-Sanchez and Wise, 2022). Resistance has even been reported against the latest RNA-interference (RNAi) based pesticide in CPB (Mishra et al., 2021). CPB is a non-model model for rapid host adaptation and insecticide resistance research (Alyokhin and Chen, 2017; Casagrande, 1987; Y. H. Chen et al., 2023). The known mechanisms of insecticide resistance in CPB are target site insensitivity, and altered detoxification, transport, penetration, and excretion of insecticides (Alyokhin et al., 2022, 2008). Reduced cholinesterase activity has been found to be involved with carbofuran resistance (Ioannidis et al., 1992; Wierenga and Hollingworth, 1994). Increased levels of monooxygenase enzymes (Rose and Brindley, 1985) and esterases (Argentine et al., 1989), have been associated with resistance against carbamate and organophosphate insecticides. Mutations typically associated with organophosphate and pyrethroid resistance, namely, the S291G of acetylcholinesterase and L1014F of sodium channel respectively, have been detected in several resistant populations (Clark et al., 2001; Lee et al., 2000; Malekmohammadi et al., 2012; Tebbe et al., 2016). Hawthorne et.al. identified three quantitative genetic loci (QTL) contributing to resistance against a pyrethroid through linkage mapping (Hawthorne, 2003).

Neonicotinoids are one of most used insecticide classes in the world (Bass et al., 2015). Imidacloprid is a neonicotinoid that acts as an inhibitor of the neurotransmitter acetylcholine causing paralysis and eventually death in insects (Bai et al., 1991). Imidacloprid has been used to control CPB without complete success for years. Imidacloprid and two other neonicotinoids are banned in some places by law, but the use of imidacloprid persists. Overexpression of a cytochrome P450 monooxygenase and a uridine diphosphate (UDP)-glycosyltransferase gene has been associated with imidacloprid resistance in CPB (Kaplanoglu et al., 2017).

To the best of our knowledge, a cross-based genetic mapping of imidacloprid resistance has not been performed on CPB before. Hence, we aimed to investigate the genetic basis of imidacloprid in CPB using the cost-effective mapping technique called bulked segregant analysis (BSA) in which individuals showing extreme phenotypes for a trait are pooled (Kurlovs et al., 2019; Li and Xu, 2022; Michelmore et al., 1991; Van Leeuwen et al., 2012; Yu et al., 2021). To this end, we generated intercross lines from five European strains of CPB and performed BSA to identify the genetic basis of resistance against imidacloprid. The key questions we ask in this study are: 1) What are the genes involved in imidacloprid resistance in CPB? 2) Are any of those genes differentially expressed in the resistant genotype in comparison to the susceptible genotype? 3) Does the knockdown of any of the differentially expressed genes affect the susceptibility levels of CPB against imidacloprid?

## 2. Materials & Methods

### 2.1. Insects, plants and insecticide

Colorado potato beetles were obtained from Dr. Ralf Nauen’s lab, Bayer AG (Monheim, Germany). The five original strains were collected by the supplier and team from different parts of Europe at different points in time (Mehlhorn et al., 2020). They are namely, D01 (Germany, 2002), E01 (Spain, 2014), E02 (Spain, 2017), E06 (Spain, 2012), and U01 (Ukraine, 2012). The E06 strain is the most resistant out of the five with a resistance ratio of 25.03 against the most susceptible D01 strain (Edison et al., 2024).

All insects had been reared on five-week-old potato (*Solanum tuberosum*) plants (Annabelle variety, seed potatoes purchased from Ellenberg’s Kartoffelvielfalt GmbH & Co. KG, Barum) under insecticide-free conditions for at least 20 generations in 85cm x 45cm x 55cm sized insect cages. The beetle rearing, plant growing, and experiments were carried out in a greenhouse (Methods sections 2.2 & 2.3) and a climate chamber (Methods section 2.7) under a long day photoperiod (16:8 L:D) at a temperature of 24°C. The detailed method of rearing is stated elsewhere (Edison et al., 2024).

Imidacloprid, a neonicotinoid insecticide, was used in our experiments in the form of water-dispersible granules available commercially by the name Confidor WG 70 (70g/Kg Imidacloprid, Bayer AG, Monheim, Germany). The insecticide granules were mixed in water and applied to the soil. Due to the discontinuation of Confidor in the EU, technical grade imidacloprid (Bayer AG, Monheim, Germany) was used for the RNAi experiments.

### 2.2. Generating the segregating population

The five strains of CPB were intercrossed according to the schematic in Figure 1A which resulted in 20 lines. For every two strains that were crossed, reciprocal crosses were made with four mating pairs in each combination. This sequential and random crossing method was used to facilitate higher resolution in mapping (Darvasi and Soller, 1995).

**Figure 1.**
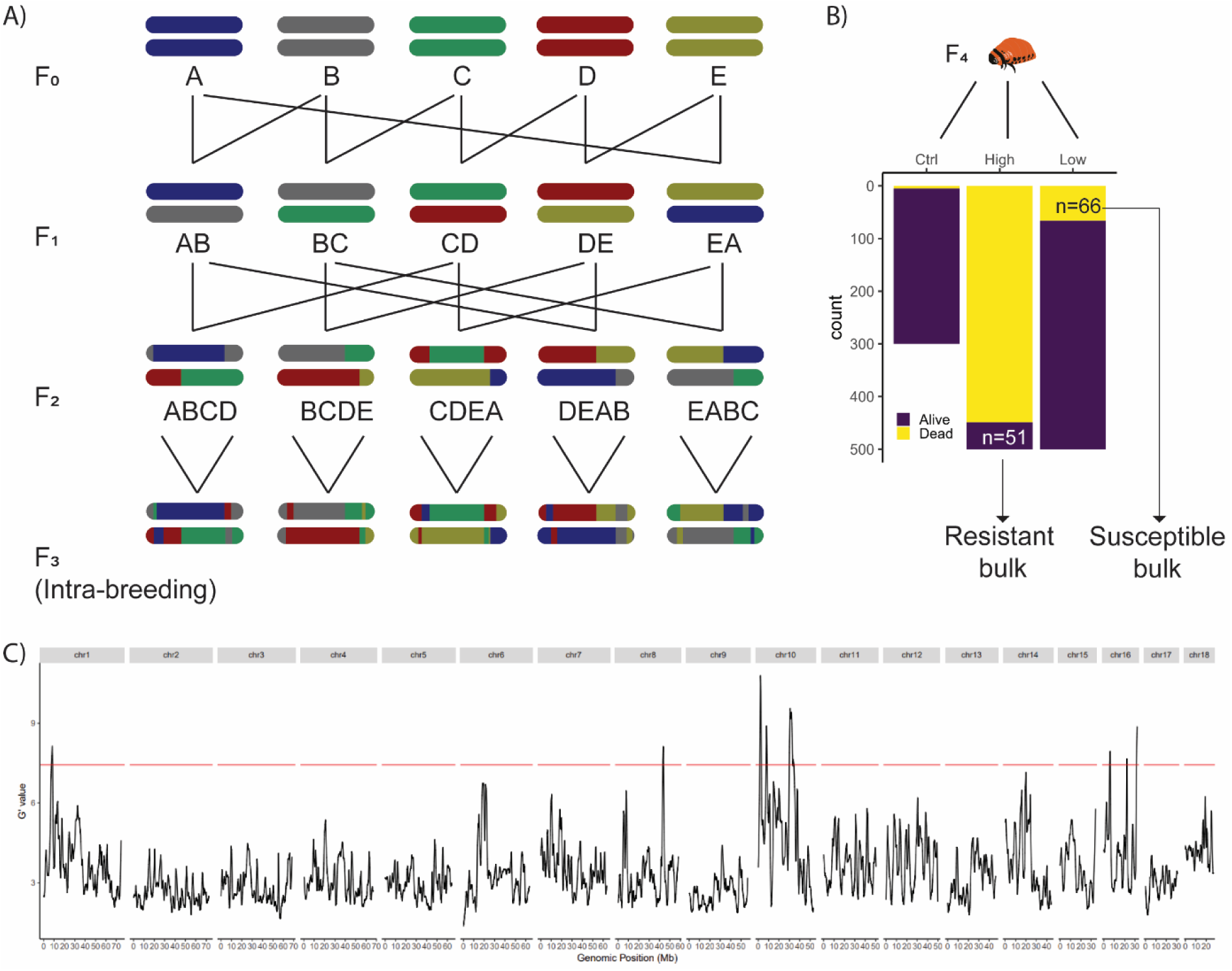
Eight QTL peaks associated with imidacloprid resistance identified through Bulk Segregant Analysis. **A. Crossing scheme.** Five strains of CPB were intercrossed to generate a genetically diverse mapping population for QTL mapping via bulked segregant analysis. The letters denote the parental strains: A= D01, B= E01, C=E02, D=E06 and E=U01. For each combination of intercrosses shown here, male x female and female x male crosses were constructed. This resulted in 10 crosses at the F1 stage and 20 crosses at the F2 stage. At the F3 stage, each cross was allowed to inbreed. **B. Phenotyping assays.** The inverted bar plot shows the number of dead (yellow) and alive (purple) larvae in the three treatments of the phenotyping assay-1) Ctrl: water 2) High: 250ppm imidacloprid 3) Low: 1ppm imidacloprid. The arrows indicate the number and type of bulk (resistant bulk: alive in the high treatment & susceptible bulk: dead in the low treatment) the larvae formed for BSA. **C. QTL peaks.** The G’ statistic is plotted for the whole genome of CPB. The chromosome number is indicated on top. The red horizontal line represents the threshold for significance. The genomic position in Mb is shown on the x-axis.

The 20 lines were allowed to mate and develop for one generation without intermixing to form the F3 generation. The offspring of this inbred generation were further subjected to phenotyping assays. While generating the crosses, for each generation, pupae were sexed (Pelletier, 1993), and the males and females were stored in separate plastic boxes. As it was crucial not to overrepresent or underrepresent each strain or cross, sex determination was once again performed on the freshly emerged adults as well. The cages and boxes were well-labeled, and insects were handled with utmost care to avoid cross-contamination.

### 2.3. Phenotyping assays

An equal number of larvae from each of the final crosses were subjected to phenotyping assays to create pools of individuals showing extreme phenotypes for imidacloprid sensitivity (highly resistant vs. highly susceptible).

Toxicity assays were performed on five-day-old F4 2^nd^ instar larvae of the segregating population to separate the individuals into two extreme bulks (Figure 1B). Insecticide was applied topically on larvae that were placed in a petri dish lined with filter paper, following the IRAC Susceptibility Test Method 029 (IRAC Methods Working Group, 2013). Two aqueous solutions containing either 1 ppm (low-concentration treatment) or 250 ppm (high-concentration) imidacloprid were used. These concentrations were chosen based on pretests so that each of the two bulks comprised of 10% of the tested individuals. In each of the two insecticide treatments, 500 larvae were tested, and 300 larvae were tested in the control treatment (water without insecticide). 1µl of the aqueous solution for the respective treatment was applied using a micropipette (control or one of the two insecticide solutions) on the top of each larva. The larvae were supplied with sufficient leaves to feed on throughout the duration of the assay to avoid the effects of starvation. Mortality was assessed after 48h, and the dead larvae (n=61) were collected from the low-concentration treatment and the surviving larvae (n=55) were collected from the high-concentration treatment to form bulks of highly susceptible and highly resistant individuals, respectively. All larvae were frozen in liquid nitrogen and stored in −20°C freezer until DNA isolation.

### 2.4. DNA isolation and Whole genome sequencing (WGS)

Genomic DNA was isolated from all pooled samples using the Monarch Genomic DNA purification kit #T3010 (New England BioLabs) following the manufacturer’s protocol. From the toxicity assays, 66 and 51 larvae were pooled to form the susceptible and the resistant bulks respectively. The concentration of DNA was assessed using Nanodrop 1000 (Thermo Scientific, Germany). Whole genome sequencing was performed using Illumina Novoseq sequencer at Novogene (Cambridge, UK).

### 2.5. Bulked segregant analysis

Read trimming was performed with Skewer (Jiang et al., 2014). Mapping was performed with BWA-*MEM* (Li, 2013) and variant calling was performed using GATK (McKenna et al., 2010). The high-quality variants were used as markers for the bulked segregant analysis using the G’ statistic (Magwene et al., 2011) implemented in the *QTLseqr* package (Mansfeld and Grumet, 2018) in R. Candidate genes were examined using the published genome annotation and the functional annotation of CPB (J. Yan et al., 2023). Using the Gene Expression Atlas (GEA) of CPB (Wilhelm et al., 2024), genes that were not sufficiently expressed were filtered out.

### 2.6. Relative expression of candidates in susceptible and resistant genotypes

Three 2^nd^ instar larvae each from the susceptible (E01) and resistant (E06) strains were used for RNA extraction, cDNA synthesis, and qPCR. Total RNA was isolated from the whole larval body using the RNeasy Mini Kit (Qiagen, Hilden, Germany) following the manufacturer’s protocol. The yield and purity were then measured using Nanodrop 1000 (Thermo Scientific, Germany). cDNA synthesis was performed with 0.1µg of total RNA using the RevertAid First-Strand cDNA synthesis kit (ThermoFisher Scientific, Germany) with oligo-dT primers. For qPCR, previously published primer sequences were used for the housekeeping genes RP18 and ARF1 (Shi et al., 2013). For the candidates, primer pairs were designed using PrimerBLAST (Ye et al., 2012). Based on a dilution series (1:10, 1:100 and 1:1000), primer efficiencies were calculated for each primer pair used. The sequences and efficiencies of the chosen primers are shown in Supplementary Table 1. Then, for performing RT-qPCR, 1:100 dilution of the cDNA was used. RT-qPCR was performed in a RotorGene Q system (Qiagen) using the KAPA SYBR FAST kit (Roche, Basel, Switzerland). The program consisted of initial denaturation at 98°C for 3 min, followed by 40 cycles of denaturation (98°C, 3s) and annealing/extension (60°C, 20s). RT-qPCR data was analyzed following the 2^−ΔΔCT^ method (Livak and Schmittgen, 2001) with expression normalized to two housekeeping genes, RP18 and ARF1.

### 2.7. Functional validation using RNAi

RNAi was performed to silence the expression of the target genes to validate their function. 2^nd^ instar larvae weighing 3.5-4.5mg were allowed to ingest leaf discs coated with 50ng/12µl double stranded RNA (dsRNA). Three types of dsRNA, namely, ds*GFP*, ds*LdNA19763*, and ds*LdNA20673* respectively targeting the enhanced Green Fluorescent Protein (eGFP) gene (GenBank: U55761.1), and the CPB genes *LdNA19763* and *LdNA20673* were designed by and purchased from Eupheria Biotech (Dresden, Germany). All dsRNA sequences were formatted using EMBL-EBI Job Dispatcher sequence analysis tools framework (Madeira et al., 2024) and are given in Supplementary Table 2. 12µl of aqueous solutions of ds*LdNA20673* (n=89) or ds*LdNA19763* (n=90) was carefully spread on a 2cm diameter leaf disc (cut from leaves at the 2^nd^-4^th^ position from the top of 5-weeks-old plants) using a pipette tip. Water (n=83) and ds*GFP* (n=88) were used as negative controls. Once the leaf discs were dry, they were placed in a petri dish lined with moistened filter paper. A single larva was placed on the leaf disc. The dsRNA treatment was repeated daily for three days with freshly treated leaf discs. After that, the larvae were fed with untreated fresh leaves for two days. On the 6^th^ day, the larvae from each treatment were assigned to two groups for toxicity assays-100ppm of technical grade imidacloprid or acetone. 2µl insecticide or acetone was applied topically on the larvae. Data were analyzed by performing a binomial generalized linear model on the counts of ‘dead’ or ‘alive’ larvae as a function of the treatment. Binomial tests of proportions were also performed to identify which proportions are different from each other. To verify the successful knockdown of the genes, three larvae were collected from each treatment and snap-frozen on the 6th day before the toxicity experiment. RT-qPCR was then performed as previously described to check the expression of the candidate genes in all the different dsRNA treatments relative to the control (water) samples. RT-qPCR data was analyzed following the 2^−ΔΔCT^ method (Livak and Schmittgen, 2001) with expression normalized to two housekeeping genes, RP18 and ARF1. The figures were created using the “ggplot2” (Wickham, 2016) and “viridis” (Garnier et al., 2023) packages. All analyses were performed using R version 4.3 (R Core Team, 2023) and RStudio version 2024.09.0.

## 3. Results

### 3.1. Eight QTL peaks governing imidacloprid resistance identified using BSA

To identify genomic regions associated with imidacloprid resistance, we first created a mapping population by intercrossing the five CPB strains following the design shown in Figure 1A. Then, toxicity assays were performed on the F4 offspring at the 2^nd^ instar larval stage with two concentrations (high and low) to yield approximately 10% of the larvae in each of the bulks (Figure 1B). After DNA isolation and whole genome sequencing, the reads were mapped to the Colorado potato beetle reference genome (J. Yan et al., 2023). After SNP calling and filtering, we used the R package QTLseqr (Mansfeld and Grumet, 2018) to calculate the G’ statistic (Magwene et al., 2011), which identified eight peaks potentially involved with imidacloprid resistance in CPB in chromosomes 1, 8, 10 and 16 (Figure 1C). In total, 171,711 SNPs were found to overlap the significant regions.

### 3.2. Candidate genes narrowed down based on the gene expression atlas and the functional annotation

Using the genome annotation of CPB, 337 genes were identified to be present in the eight peaks. A full list of those 337 genes along with information about their potential function is given in Supplementary Table 2. Using the GEA of CPB (Wilhelm et al., 2024), 126 genes were found to be sufficiently expressed (>5 Transcripts Per Million (TPM)) in the first, second or third instar larvae. Among them, SNPs predicted to result in at least one nonsynonymous amino acid change were identified to be present in 65 genes using SNPEff (Cingolani et al., 2012) (Table 1, Supplementary Figure 1). Then, using the functional annotation of CPB, we manually searched for genes that are functionally commonly associated with resistance including cytochrome P450 monooxygenases (CYPs), ATP-binding cassette (ABC) transporters and glutathione S-transferases (GSTs). Two ABC transporters (*LdNA19763*, *LdNA19944*) and one galactosyl transferase (*LdNA20673*) were found on the 10^th^ chromosome. The expression profiles of these three genes across the first, second and third larval instars were downloaded from the GEA website (Wilhelm et al., 2024) (Supplementary Figure 2).

**Table 1.**
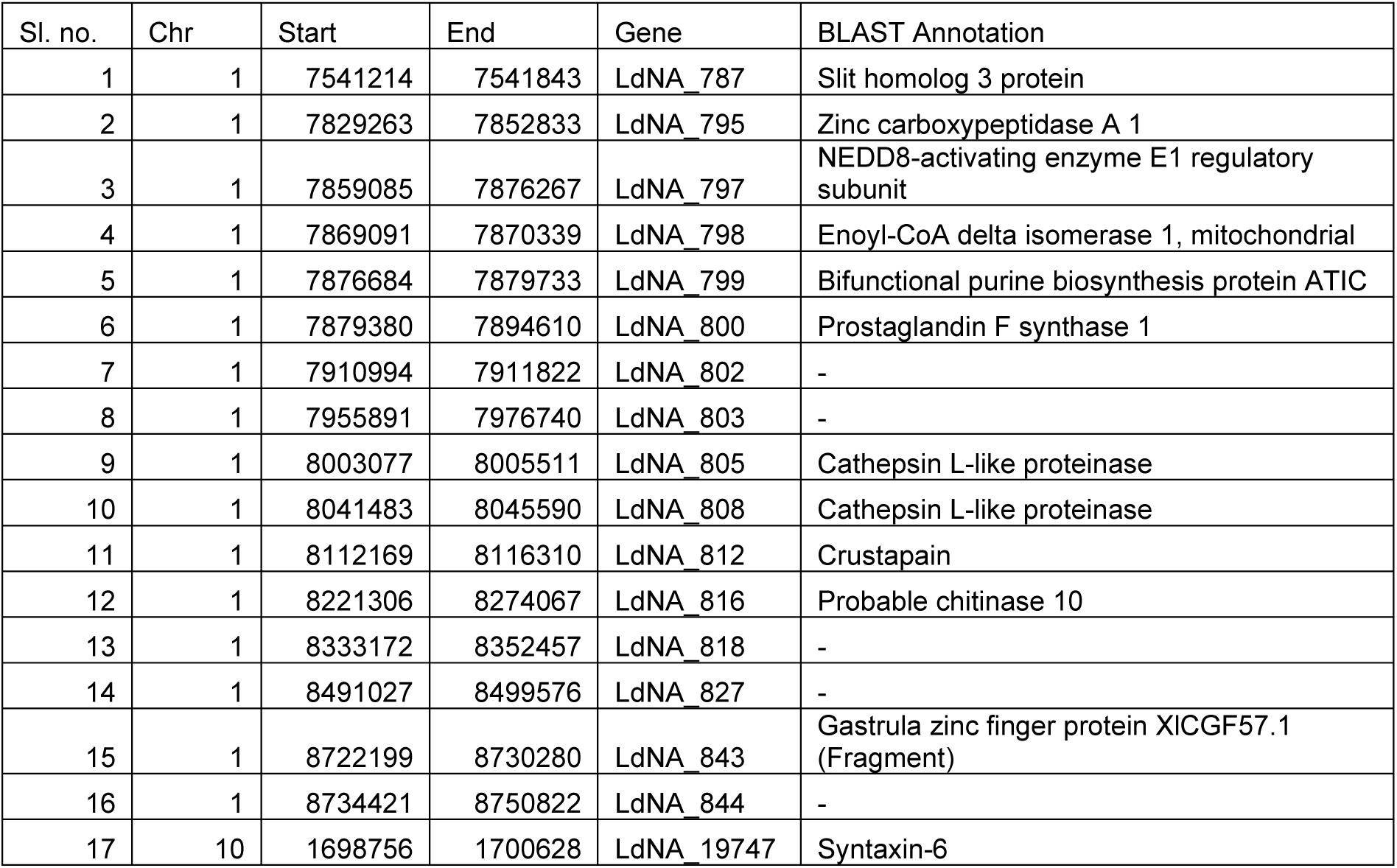
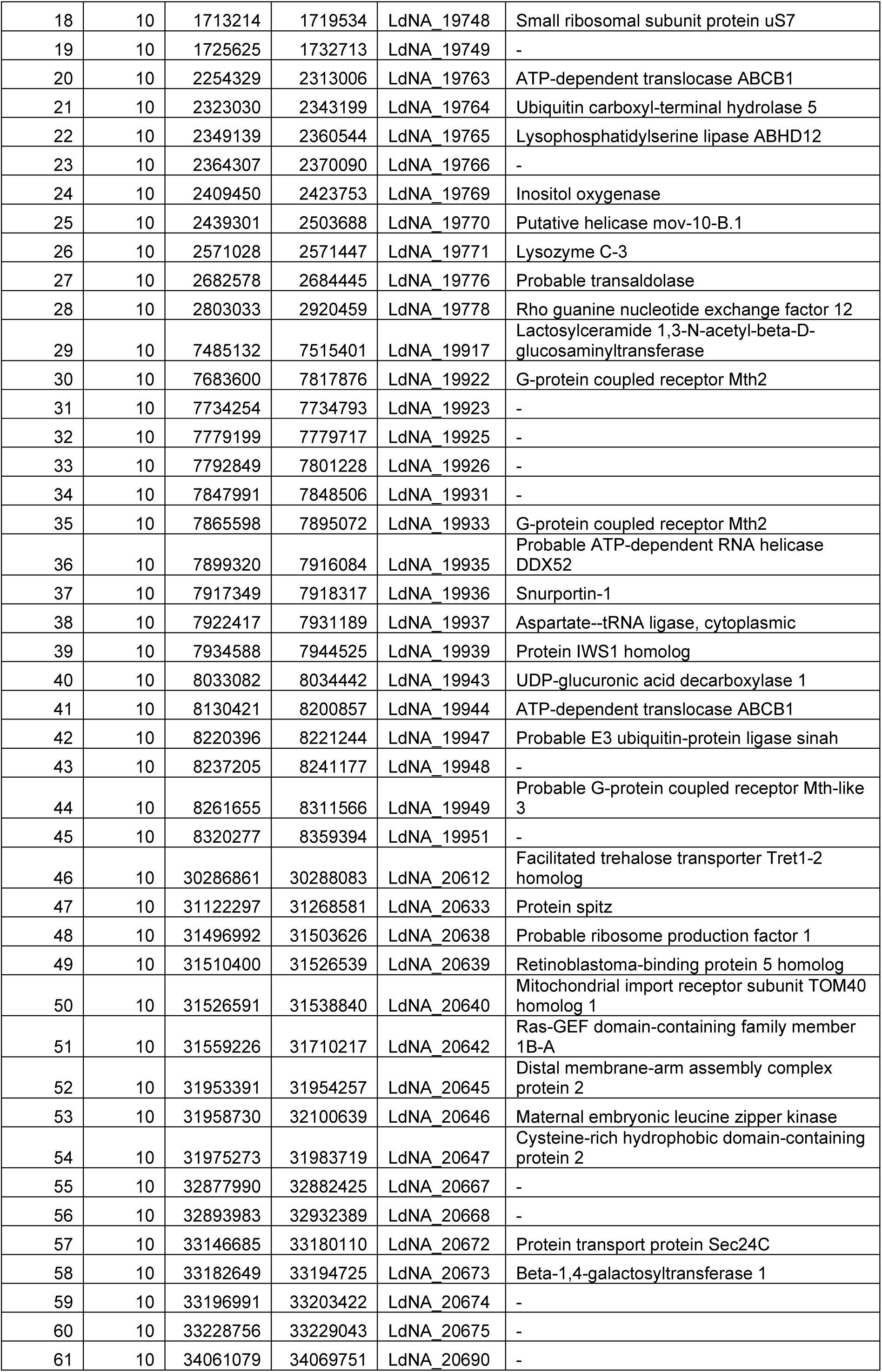
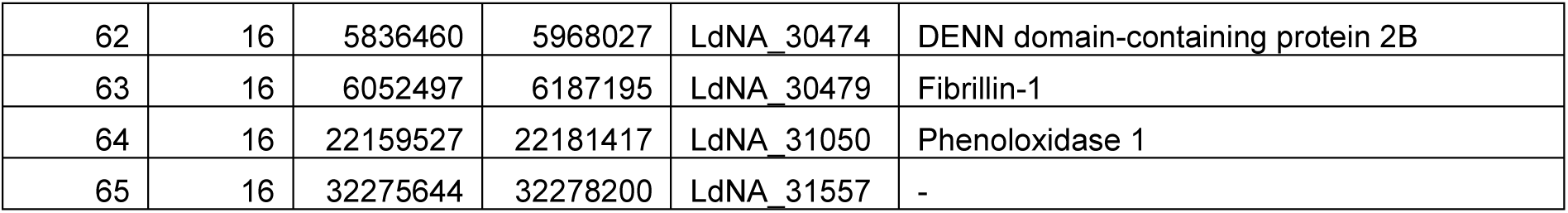
List of 65 candidate genes. A list of genes that are sufficiently expressed in at least one larval stage of CPB (Wilhelm et al., 2024) and additionally contain SNPs that result in at least one amino acid change. The potential function of the gene is given in the ‘BLAST Annotation’ column if available; ‘-’ indicates lack of information. The ‘Start’ and ‘End’ columns indicate the position of the gene in bp within the specific chromosome.

### 3.3. Relative expression of candidates in the susceptible and resistant strains

We quantified the expression differences of the candidate genes between a resistant and a susceptible strain. Two out of the three candidates (*LdNA19763* and *LdNA20673*) are expressed in higher amounts in the relatively more susceptible E01 strain compared to the relatively more resistant E06 strain (*LdNA19763*: p<0.001, *LdNA19944*: p=0.387, *LdNA20673*: p=0.055, t-test, Figure 2). The expression of *LdNA19763* and *LdNA20673* are elevated in the E01 strain approximately by factors of 3.6 and 2.2 respectively.

**Figure 2.**
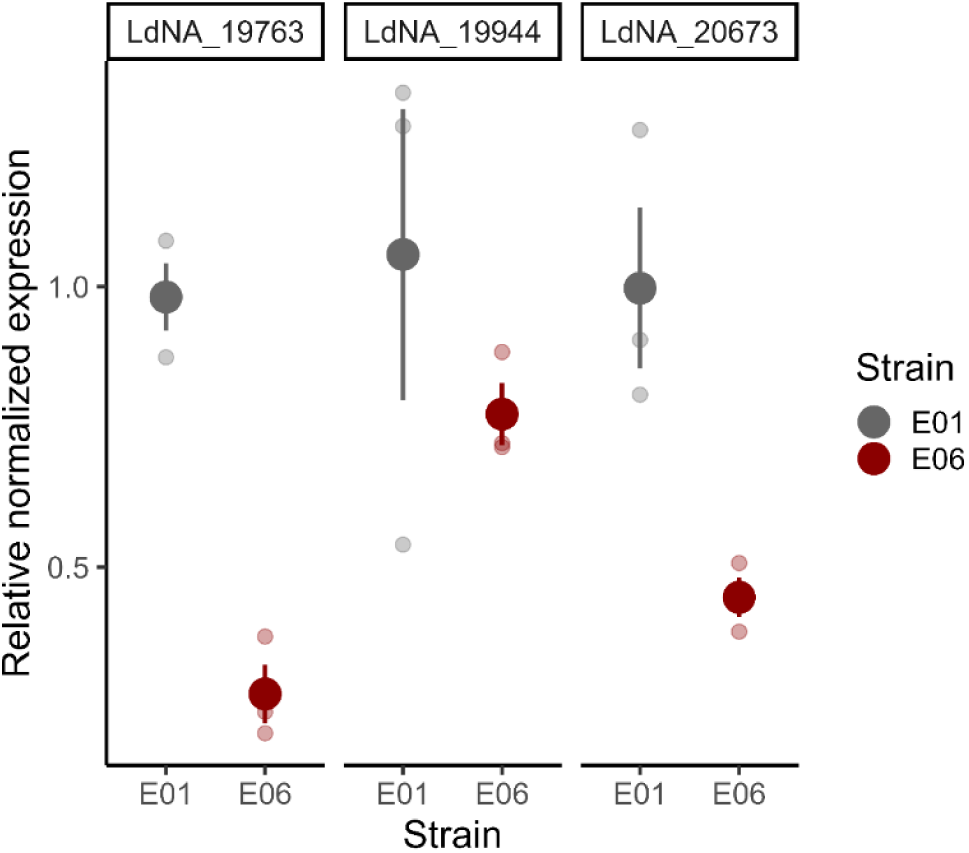
Relative expression of three candidate genes in two strains. The graph shows the relative expression of three candidate genes (LdNA_19763, LdNA_19944 and LdNA_20673). The colors indicate the two strains (Grey: E01 and Red: E06). The expression was calculated relative to the comparatively more susceptible strain, E01. The opaque dots show the mean values while the smaller faded dots show the value for the three replicates. The error bars show the standard error of the mean.

### 3.4. Functional validation of the candidates

To functionally validate the two differentially expressed candidates, RNAi was performed for each gene followed by toxicity experiments. Due to the unavailability of the susceptible E01 strain, knockdowns were performed on the resistant E06 strain. To verify the knockdown of the candidates, RT-qPCR was performed. The expression data shows high variation, and the knockdowns were partially successful (Figure 3B). Expression of gene *LdNA20673* differed significantly between dsGFP and ds*LdNA20673* treatments (p=0.03, t-test). Expression of gene *LdNA19763* differed between ds*LdNA19763* and ds*LdNA20673* treatments (p=0.04, t-test).

**Figure 3.**
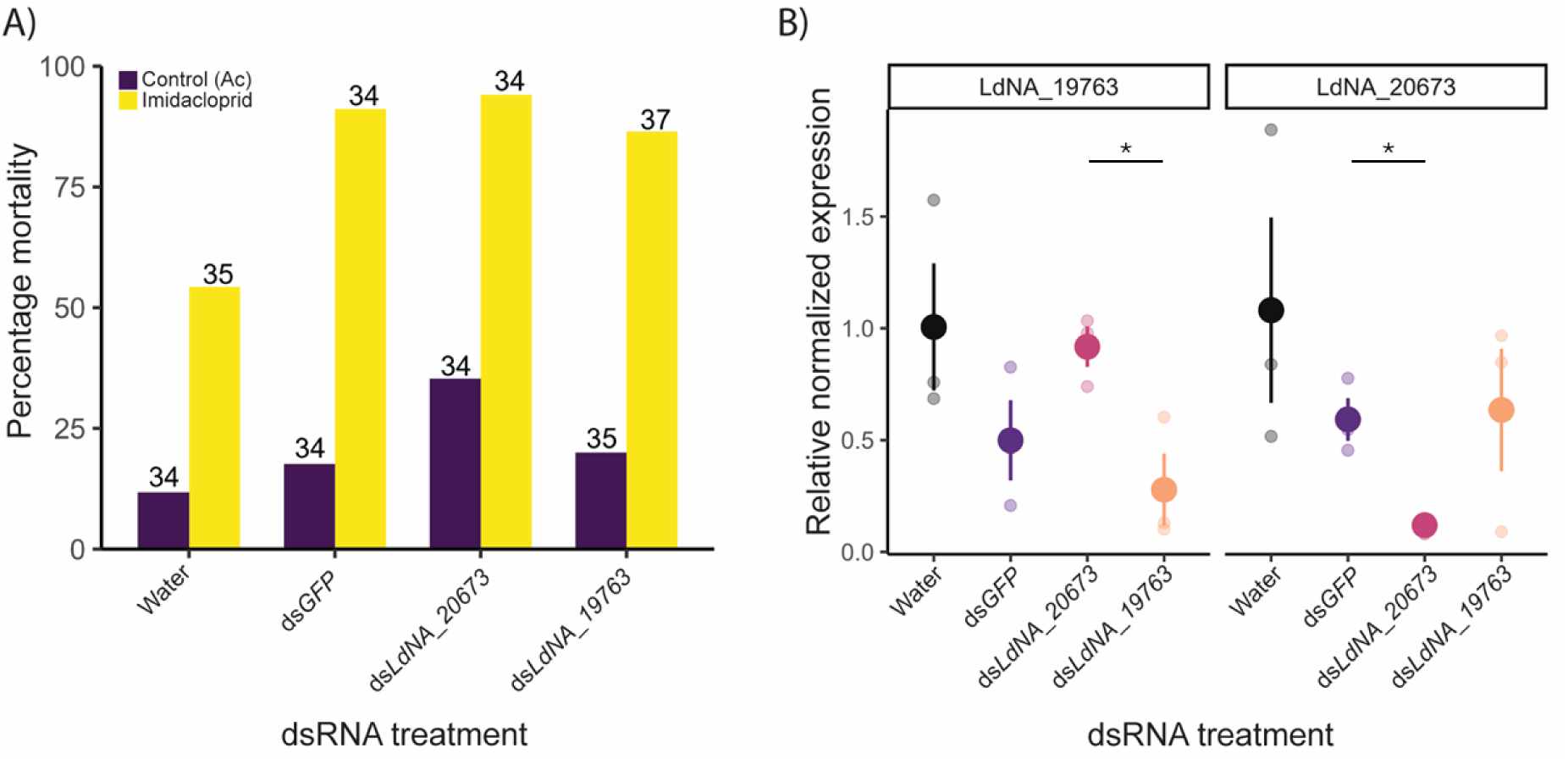
Functional validation of two candidates using RNAi. **A. Mortality after insecticide treatment.** The bar plot shows the percentage mortality after insecticide and control treatments for each of the different dsRNA treatment groups. (*LdNA19763*, *LdNA19944* and *LdNA20673*). The colors indicate the two treatments in the toxicity assay (Purple: Control (acetone) and Yellow: Imidacloprid). The numbers above the bars are the number of larvae tested in each group. **B. Verification of knockdown after RNAi.** The graph shows the relative expression of the two knocked-down genes (*LdNA19763* and *LdNA20673*) in the different dsRNA treatments (x-axis). Asterisks indicate significant differences in expression. The expression was calculated relative to the water control. The opaque dots show the mean values while the smaller faded dots show the value for the three replicates. The error bars show the standard error of the mean.

The final mortality is affected by the dsRNA treatment type (p<0.001, GLM: Status∼Treatment, family= binomial, Figure 3A). Mortality due to the insecticide treatment is significantly lower in the water treatment than in the other treatments (p<0.001, binomial test of proportions) while the control (acetone) mortality is not different between the treatments (p=0.1, binomial test of proportions). The mortality after knockdown of genes *LdNA19763* or *LdNA20673* is not different from that in the ds*GFP* treatment (p=0.5, binomial test of proportions). Due to the difference between the two negative controls, we do not find any conclusive evidence for the involvement of these two genes in resistance against imidacloprid in CPB.

## 4. Discussion

In this study, we attempted to identify the genetic basis of resistance against imidacloprid in the Colorado potato beetle by intercrossing five strains of CPB and performing bulked segregant analysis. We found eight QTL peaks associated with resistance in which an ABC transporter gene and a galactosyl transferase gene were expressed in higher amounts in the susceptible strain when compared to the resistant strain. We then performed functional validation of the two genes using RNAi. We did not find conclusive evidence for the sole involvement of the two genes in resistance to imidacloprid in CPB, largely due to methodological limitations.

While manually searching for genes that have some functional relevance in resistance, we found three ABC transporters belonging to the B subfamily in the peaks. The role of ABC transporters in insecticide resistance in CPB is not as clear as in other insects like *Drosophila* or *Tribolium*. Due to their natural excretory functions and the similarity of many insecticides to the substrates of ABC transporters, they have been implicated in insecticide resistance (Amezian et al., 2024; Dermauw and Van Leeuwen, 2014; Gott et al., 2017). In support of this, the knockdown of some ABC transporters has been found to affect the susceptibility of CPB to imidacloprid (Gaddelapati et al., 2018) and Cry toxins (Güney et al., 2021). At the same time, some studies have found that the knockdown of certain ABC transporters does not affect susceptibility levels of CPB against imidacloprid (Kaplanoglu et al., 2017) or ivermectin (Favell et al., 2020). Further studies on the molecular functions of the ABC transporters in CPB might help understand the detailed mechanism behind the involvement of these enzymes in insecticide transport and resistance.

Numerous previous studies have shown the involvement of CYP genes across several insect species (Daborn et al., 2002; Kaplanoglu et al., 2017; Nauen et al., 2022; Zhu et al., 2016). Surprisingly, no CYPs were found among the 337 genes identified here (Supplementary Table 3). CPB possesses about 74-98 CYP genes that carry out different functions including the detoxification of insecticides (Nauen et al., 2022; Wan et al., 2013). Changes in the CYP genes may constitutively increase their expression in resistant populations of several insect species, thus contributing to increased resistance. Additionally, transcription factors like the cap ‘n’ collar isoform C (CncC) have also been shown to regulate the increased expression of CYPs and the subsequent detoxification of insecticides (Gaddelapati et al., 2018; Kalsi and Palli, 2017). Lv et al. have recently demonstrated that the CF2-II transcription factor negatively regulates the expression of ABC transporters and influence insecticide resistance in cotton aphids (Lv et al., 2023). Interestingly, we found nine C2H2 type zinc finger genes in our peaks (Supplementary Table 3).

Reduced expression of genes associated with insecticide resistance in resistant strains compared to susceptible strains is inconsistent with the general understanding of resistance mechanisms. However, such a pattern is not unheard of for many resistance-associated genes, including ABCs and CYPs (Dively et al., 2020; Guo et al., 2015; Kaplanoglu et al., 2017; Zhu et al., 2016). It is unclear how such an expression pattern might be regulated in resistant populations. Negative regulatory pathways mediated by transcription factors like CnCC or C2H2 zinc finger proteins may be involved (Lv et al., 2023; Palli, 2020).

Other potential candidates include five chemoreceptors that we found in the peak in chromosome 1. Chemoreceptor genes have been found to be overexpressed in some resistant insect populations (Li et al., 2023; Xu et al., 2022). Chemosensory proteins and other sensory receptor genes were initially implicated in resistance due to their potential role in altering behavior. Additionally, due to their versatility in transporting and sequestering several types of compounds, chemosensory proteins are also garnering attention as a novel mechanism of resistance (Pelosi et al., 2018; Pu and Chung, 2024).

Advances in functional genomic techniques and the vast amount of genomic data have made it possible to investigate the role of specific genes in insecticide resistance in many non-model pests (Homem and Davies, 2018; Przybyla and Gilbert, 2022). In our study, we attempted to functionally validate the role of two genes, namely, *LdNA19763* and *LdNA20673*. We found no differences in the mortalities after insecticide treatment among the different dsRNA treatments. There was a significant difference between the two controls (ds*GFP* and water) upon imidacloprid treatment. Mortality was higher in the ds*GFP* treatment than in the water treatment while there was no difference in the control (acetone) mortality. This is intriguing, since we found no clear potential target of ds*GFP* in the CPB genome. Differences in mortality between the two controls in RNAi treatments (water and ds*GFP*), although not significantly large, have been observed in other studies (Kishk et al., 2017; Shi et al., 2023). However, it is unclear whether this is simply stochastic or if the ds*GFP* treatment is indeed associated with increased mortality. Higher mortality observed in the ds*GFP* treatment could be due to toxicity caused by the dsRNA molecule itself or any off-target effects on important genes. There is some evidence for ds*GFP* affecting the expression of nontarget genes in honeybees (Jarosch and Moritz, 2012; Nunes et al., 2013). Since the control (acetone) mortality is not significantly higher in the ds*GFP* treatment compared to the water treatment in our experiments, it is more plausible that the ds*GFP* treatment enhances the stress caused by the insecticide. Due to this methodological limitation, our RNAi experiments did not provide clear support on the role of the two candidates in imidacloprid resistance. Our study raised an important concern on the suitability of using RNAi to validate genetic mechanisms underlying stress resistance in CPB.

## Acknowledgements

We thank Ursula Martiné for DNA isolation, subsequent preparation of samples for sequencing and highly valuable tips for working in the molecular lab. We are grateful to Thilo Fahlbusch for helping maintain crosses and carry out toxicity assays. We also thank Léonore Wilhelm and Yangzi Wang for early access to the CPB Gene Expression Atlas and the CPB annotations respectively. We further thank Dirk Schmidt and Sascha Ahrens for their assistance in growing potato plants in the greenhouse. We also thank Holger Schön and David Martin Fernandez for constructing the insect cages.

## Data availability statement

All genomic sequences used in this study are deposited in NCBI under the Bioproject PRJNA1121994.

## Author contributions

SX conceived, designed, and supervised the study. AE and NN designed and conducted the experiments. PD performed all bioinformatic analyses including mapping, variant calling, BSA, and SNP annotation. AE analyzed the experimental data. SX acquired funding for this research. AE wrote the paper. NN, PD and SX reviewed and revised the manuscript substantially. All authors read and approved the manuscript for submission.

## References

Alyokhin, A., Chen, Y.H., 2017. Adaptation to toxic hosts as a factor in the evolution of insecticide resistance. Curr Opin Insect Sci. 10.1016/j.cois.2017.04.006

Alyokhin, A., Rondon, S.I., Gao, Y., 2022. Insect pests of potato, Insect Pests of Potato. Elsevier.

Alyokhin, Andrei, Baker, Mitchell, Mota-Sanchez, David, Dively, Galen, Grafius, Edward, Alyokhin, A, Baker, M, Mota-Sanchez, D, Grafius, E, Dively, G, 2008. Colorado Potato Beetle Resistance to Insecticides. J. Pot Res 85, 395–413. 10.1007/s12230-008-9052-0

Amezian, D., Nauen, R., Van Leeuwen, T., 2024. The role of ATP-binding cassette transporters in arthropod pesticide toxicity and resistance. Curr Opin Insect Sci. 10.1016/j.cois.2024.101200

Argentine, J.A., Clark, M.J., Ferro, D.N., 1989. Genetics and Synergism of Resistance to Azinphosmethyl and Permethrin in the Colorado Potato Beetle (Coleoptera: Chrysomelidae). J Econ Entomol 82, 698–705. 10.1093/jee/82.3.698

Bai, D., Lummis, S.C.R., Leicht, W., Breer, H., Sattelle, D.B., 1991. Actions of imidacloprid and a related nitromethylene on cholinergic receptors of an identified insect motor neurone. Pestic Sci 33, 197–204. 10.1002/ps.2780330208

Bass, C., Denholm, I., Williamson, M.S., Nauen, R., 2015. The global status of insect resistance to neonicotinoid insecticides. Pestic Biochem Physiol. 10.1016/j.pestbp.2015.04.004

Casagrande, R.A., 1987. The Colorado potato beetle: 125 years of mismanagement. Bulletin of the Entomological Society of America 33, 142–150. 10.1093/besa/33.3.142

Chen, L.P., Jiang, H.Q., Luo, L., Qiu, J., Xing, X.J., Hou, R.Y., Wu, Y.J., 2023. The role of intercellular junction proteins in the penetration resistance of *Drosophila* larvae to avermectin. Comparative Biochemistry and Physiology Part - C: Toxicology and Pharmacology 266. 10.1016/j.cbpc.2023.109557

Chen, Y.H., Cohen, Z.P., Bueno, E.M., Christensen, B.M., Schoville, S.D., 2023. Rapid evolution of insecticide resistance in the Colorado potato beetle, *Leptinotarsa decemlineata*. Curr Opin Insect Sci 55, 101000. 10.1016/j.cois.2022.101000

Cingolani, P., Platts, A., Wang, L.L., Coon, M., Nguyen, T., Wang, L., Land, S.J., Lu, X., Ruden, D.M., 2012. A program for annotating and predicting the effects of single nucleotide polymorphisms, SnpEff. Fly (Austin) 6, 80–92. 10.4161/fly.19695

Clark, J.M., Lee, S.H., Kim, H.J., Yoon, K.S., Zhang, A., 2001. DNA-based genotyping techniques for the detection of point mutations associated with insecticide resistance in Colorado potato beetle *Leptinotarsa decemlineata*. Pest Manag Sci 57, 968–974. 10.1002/ps.369

Daborn, P.J., Yen, J.L., Bogwitz, M.R., Le Goff, G., Feil, E., Jeffers, S., Tijet, N., Perry, T., Heckel, D., Batterham, P., Feyereisen, R., Wilson, T.G., ffrench-Constant, R.H., 2002. A Single P450 Allele Associated with Insecticide Resistance in *Drosophila*. Science (1979) 297, 2253–2256. 10.1126/science.1074170

Darvasi, A., Soller, M., 1995. Advanced intercross lines, an experimental population for fine genetic mapping. Genetics 141, 1199–1207. 10.1093/genetics/141.3.1199

Dermauw, W., Van Leeuwen, T., 2014. The ABC gene family in arthropods: Comparative genomics and role in insecticide transport and resistance. Insect Biochem Mol Biol 45, 89–110. 10.1016/j.ibmb.2013.11.001

Dively, G.P., Crossley, M.S., Schoville, S.D., Steinhauer, N., Hawthorne, D.J., 2020. Regional differences in gene regulation may underlie patterns of sensitivity to novel insecticides in *Leptinotarsa decemlineata*. Pest Manag Sci 76, 4278–4285. 10.1002/ps.5992

Edison, A., Michelbach, A., Sowade, D., Kertzel, H., Schmidt, L., Schäfer, M., Lysander, M., Nauen, R., Duchen, P., Xu, S., 2024. Evidence of active oviposition avoidance to systemically applied imidacloprid in the Colorado potato beetle. Insect Sci 0, 1–12. 10.1111/1744-7917.13319

Enayati, A.A., Ranson, H., Hemingway, J., 2005. Insect glutathione transferases and insecticide resistance. Insect Mol Biol. 10.1111/j.1365-2583.2004.00529.x

Favell, G., McNeil, J.N., Donly, C., 2020. The ABCB Multidrug Resistance Proteins Do Not Contribute to Ivermectin Detoxification in the Colorado Potato Beetle, *Leptinotarsa decemlineata* (Say). Insects 11, 135. 10.3390/insects11020135

ffrench-Constant, R.H., Rocheleau, T.A., Steichen, J.C., Chalmers, A.E., 1993. A point mutation in a Drosophila GABA receptor confers insecticide resistance. Nature 363, 449–451. 10.1038/363449a0

Gaddelapati, S.C., Kalsi, M., Roy, A., Palli, S.R., 2018. Cap ‘n’ collar C regulates genes responsible for imidacloprid resistance in the Colorado potato beetle, *Leptinotarsa decemlineata*. Insect Biochem Mol Biol 99, 54–62. 10.1016/j.ibmb.2018.05.006

Garnier, S., Ross, N., Rudis, boB, Filipovic-Pierucci, A., Galili, T., timelyportfolio, O’Callaghan, A., Greenwell, B., Sievert, C., Harris, D.J., Sciaini, M., Chen, J., 2023. sjmgarnier/viridis: CRAN release v0.6.3. 10.5281/zenodo.7890878

Gott, R.C., Kunkel, G.R., Zobel, E.S., Lovett, B.R., Hawthorne, D.J., 2017. Implicating ABC transporters in insecticide resistance: Research strategies and a decision framework. J Econ Entomol 110, 667–677. 10.1093/jee/tox041

Güney, G., Cedden, D., Hänniger, S., Heckel, D.G., Coutu, C., Hegedus, D.D., Mutlu, D.A., Suludere, Z., Sezen, K., Güney, E., Toprak, U., 2021. Silencing of an ABC transporter, but not a cadherin, decreases the susceptibility of Colorado potato beetle larvae to *Bacillus thuringiensis* ssp. *tenebrionis* Cry3Aa toxin. Arch Insect Biochem Physiol 108. 10.1002/arch.21834

Guo, Z., Kang, S., Zhu, X., Xia, J., Wu, Q., Wang, S., Xie, W., Zhang, Y., 2015. Down-regulation of a novel ABC transporter gene (Pxwhite) is associated with Cry1Ac resistance in the diamondback moth, *Plutella xylostella* (L.). Insect Biochem Mol Biol 59, 30–40. 10.1016/j.ibmb.2015.01.009

Hawkins, N.J., Bass, C., Dixon, A., Neve, P., 2019. The evolutionary origins of pesticide resistance. Biological Reviews 94, 135–155. 10.1111/brv.12440

Hawthorne, D.J., 2003. Quantitative trait locus mapping of pyrethroid resistance in Colorado potato beetle, *Leptinotarsa decemlineata* (Say) (Coleoptera: Chrysomelidae). J Econ Entomol 96, 1021–1030. 10.1603/0022-0493-96.4.1021

Hemingway, J., 2000. The molecular basis of two contrasting metabolic mechanisms of insecticide resistance. Insect Biochem Mol Biol 30, 1009–1015. 10.1016/S0965-1748(00)00079-5

Homem, R.A., Davies, T.G.E., 2018. An overview of functional genomic tools in deciphering insecticide resistance. Curr Opin Insect Sci 27, 103–110. 10.1016/j.cois.2018.04.004

Ioannidis, P.M., Grafius, E.J., Wierenga, J.M., Whalon, M.E., Hollingworth, R.M., 1992. Selection, inheritance and characterization of carbofuran resistance in the Colorado potato beetle (Coleoptera: Chrysomelidae). Pestic Sci 35, 215–222. 10.1002/ps.2780350304

IRAC Methods Working Group, 2013. https://irac-online.org/content/uploads/Method_029_Stinkbugs.pdf.

Jarosch, A., Moritz, R.F.A., 2012. RNA interference in honeybees: Off-target effects caused by dsRNA. Apidologie 43, 128–138. 10.1007/s13592-011-0092-y

Jiang, H., Lei, R., Ding, S.-W., Zhu, S., 2014. Skewer: a fast and accurate adapter trimmer for next-generation sequencing paired-end reads. BMC Bioinformatics 15, 182. 10.1186/1471-2105-15-182

Kalsi, M., Palli, S.R., 2017. Transcription factor cap n collar C regulates multiple cytochrome P450 genes conferring adaptation to potato plant allelochemicals and resistance to imidacloprid in *Leptinotarsa decemlineata* (Say). Insect Biochem Mol Biol 83, 1–12. 10.1016/j.ibmb.2017.02.002

Kaplanoglu, E., Chapman, P., Scott, I.M., Donly, C., 2017. Overexpression of a cytochrome P450 and a UDP-glycosyltransferase is associated with imidacloprid resistance in the Colorado potato beetle, *Leptinotarsa decemlineata*. Sci Rep 7, 1762. 10.1038/s41598-017-01961-4

Kishk, A., Anber, H.A.I., AbdEl-Raof, T.K., El-Sherbeni, A.E.H.D., Hamed, S., Gowda, S., Killiny, N., 2017. RNA interference of carboxyesterases causes nymph mortality in the Asian citrus psyllid, *Diaphorina citri*. Arch Insect Biochem Physiol 94. 10.1002/arch.21377

Kurlovs, A.H., Snoeck, S., Kosterlitz, O., Van Leeuwen, T., Clark, R.M., 2019. Trait mapping in diverse arthropods by bulked segregant analysis. Curr Opin Insect Sci 36, 57–65. 10.1016/j.cois.2019.08.004

Lee, S.H., Yoon, K.-S., Williamson, M.S., Goodson, S.J., Takano-Lee, M., Edman, J.D., Devonshire, A.L., Marshall Clark, J., 2000. Molecular analysis of kdr-like resistance in permethrin-resistant strains of head lice, *Pediculus capitis*. Pestic Biochem Physiol 66, 130–143. 10.1006/pest.1999.2460

Li, H., 2013. Aligning sequence reads, clone sequences and assembly contigs with BWA-MEM. ArXiv 1303.3997. 10.6084/M9.FIGSHARE.963153.V1

Li, X., Liu, X., Lu, W., Yin, X., An, S., 2022. Application progress of plant-mediated RNAi in pest control. Front Bioeng Biotechnol 10. 10.3389/fbioe.2022.963026

Li, Y., Ni, S., Wang, Y., Li, R., Sun, H., Ye, X., Tian, Z., Zhang, Y., Liu, J., 2023. The chemosensory protein 1 contributes to indoxacarb resistance in *Plutella xylostella* (L.). Pest Manag Sci 79, 2456–2468. 10.1002/ps.7415

Li, Z., Xu, Y., 2022. Bulk segregation analysis in the NGS era: a review of its teenage years. The Plant Journal 109, 1355–1374. 10.1111/tpj.15646

Liu, N., Li, M., Gong, Y., Liu, F., Li, T., 2015. Cytochrome P450s - Their expression, regulation, and role in insecticide resistance. Pestic Biochem Physiol 120, 77–81. 10.1016/j.pestbp.2015.01.006

Livak, K.J., Schmittgen, T.D., 2001. Analysis of relative gene expression data using real-time quantitative PCR and the 2-ΔΔCT method. Methods 25, 402–408. 10.1006/meth.2001.1262

Lv, Y., Pan, Y., Li, J., Ding, Y., Yu, Z., Yan, K., Shang, Q., 2023. The C2H2 zinc finger transcription factor CF2-II regulates multi-insecticide resistance-related gut-predominant ABC transporters in *Aphis gossypii* Glover. Int J Biol Macromol 253. 10.1016/j.ijbiomac.2023.126765

Madeira, F., Madhusoodanan, N., Lee, J., Eusebi, A., Niewielska, A., Tivey, A.R.N., Lopez, R., Butcher, S., 2024. The EMBL-EBI Job Dispatcher sequence analysis tools framework in 2024. Nucleic Acids Res 52, W521–W525. 10.1093/nar/gkae241

Magwene, P.M., Willis, J.H., Kelly, J.K., 2011. The statistics of bulk segregant analysis using next generation sequencing. PLoS Comput Biol 7. 10.1371/JOURNAL.PCBI.1002255

Malekmohammadi, M., Hejazi, M.J., Mossadegh, M.S., Galehdari, H., Khanjani, M., Goodarzi, M.T., 2012. Molecular diagnostic for detecting the acetylcholinesterase mutations in insecticide-resistant populations of Colorado potato beetle, *Leptinotarsa decemlineata* (Say). Pestic Biochem Physiol 104, 150–156. 10.1016/j.pestbp.2012.06.004

Mansfeld, B.N., Grumet, R., 2018. QTLseqr: An R Package for Bulk Segregant Analysis with Next-Generation Sequencing. Plant Genome 11. 10.3835/plantgenome2018.01.0006

McKenna, A., Hanna, M., Banks, E., Sivachenko, A., Cibulskis, K., Kernytsky, A., Garimella, K., Altshuler, D., Gabriel, S., Daly, M., DePristo, M.A., 2010. The genome analysis toolkit: A MapReduce framework for analyzing next-generation DNA sequencing data. Genome Res 20, 1297–1303. 10.1101/gr.107524.110

Mehlhorn, S.G., Geibel, S., Bucher, G., Nauen, R., 2020. Profiling of RNAi sensitivity after foliar dsRNA exposure in different European populations of Colorado potato beetle reveals a robust response with minor variability. Pestic Biochem Physiol 166, 104569. 10.1016/j.pestbp.2020.104569

Michelmore, R.W., Paran, I., Kesseli, R. V, 1991. Identification of markers linked to disease-resistance genes by bulked segregant analysis: a rapid method to detect markers in specific genomic regions by using segregating populations. Proceedings of the National Academy of Sciences 88, 9828–9832. 10.1073/pnas.88.21.9828

Mishra, S., Dee, J., Moar, W., Dufner-Beattie, J., Baum, J., Dias, N.P., Alyokhin, A., Buzza, A., Rondon, S.I., Clough, M., Menasha, S., Groves, R., Clements, J., Ostlie, K., Felton, G., Waters, T., Snyder, W.E., Jurat-Fuentes, J.L., 2021. Selection for high levels of resistance to double-stranded RNA (dsRNA) in Colorado potato beetle (*Leptinotarsa decemlineata* Say) using non-transgenic foliar delivery. Sci Rep 11. 10.1038/s41598-021-85876-1

Mota-Sanchez, D., Wise, J.C., 2022. The Arthropod Pesticide Resistance Database. Michigan State University.

Mutero, A., Pralavorio, M., Bride, J.M., Fournier, D., 1994. Resistance-associated point mutations in insecticide-insensitive acetylcholinesterase. Proceedings of the National Academy of Sciences 91, 5922–5926. 10.1073/pnas.91.13.5922

Nauen, R., Bass, C., Feyereisen, R., Vontas, J., 2022. The Role of Cytochrome P450s in Insect Toxicology and Resistance. Annu Rev Entomol 67, 105–124. 10.1146/annurev-ento-070621-061328

Nunes, F.M.F., Aleixo, A.C., Barchuk, A.R., Bomtorin, A.D., Grozinger, C.M., Simões, Z.L.P., 2013. Non-target effects of green fluorescent protein (GFP)-derived double-stranded RNA (dsRNA-GFP) used in honey bee RNA interference (RNAi) assays. Insects 4, 90–103. 10.3390/insects4010090

Palli, S.R., 2020. CncC/Maf-mediated xenobiotic response pathway in insects. Arch Insect Biochem Physiol 104. 10.1002/arch.21674

Palli, S.R., 2014. RNA interference in Colorado potato beetle: steps toward development of dsRNA as a commercial insecticide. Curr Opin Insect Sci 6, 1–8. 10.1016/j.cois.2014.09.011

Pavlidi, N., Vontas, J., Van Leeuwen, T., 2018. The role of glutathione S-transferases (GSTs) in insecticide resistance in crop pests and disease vectors. Curr Opin Insect Sci 27, 97–102. 10.1016/j.cois.2018.04.007

Pelletier, Y., 1993. A method for sex determination of the Colorado potato beetle pupa, Leptinotarsa decemlineata (Coleoptera: Chrysomelidae) 140–142.

Pelosi, P., Iovinella, I., Zhu, J., Wang, G., Dani, F.R., 2018. Beyond chemoreception: diverse tasks of soluble olfactory proteins in insects. Biological Reviews 93, 184–200. 10.1111/brv.12339

Przybyla, L., Gilbert, L.A., 2022. A new era in functional genomics screens. Nat Rev Genet 23, 89–103. 10.1038/s41576-021-00409-w

Pu, J., Chung, H., 2024. New and emerging mechanisms of insecticide resistance. Curr Opin Insect Sci 63, 101184. 10.1016/j.cois.2024.101184

R Core Team, 2023. R: a language and environment for statistical computing.

Rose, R.L., Brindley, W.A., 1985. An evaluation of the role of oxidative enzymes in Colorado potato beetle resistance to carbamate insecticides. Pestic Biochem Physiol 23, 74–84. 10.1016/0048-3575(85)90080-X

Shi, C., Tian, Y., Wang, Y., Guo, W., Jiang, W., 2023. The interaction of nicotinic acetylcholine receptor subunits Ldα3, Ldα8 and Ldβ1 with neonicotinoids in Colorado potato beetle, *Leptinotarsa decemlineata*. Pestic Biochem Physiol 195. 10.1016/j.pestbp.2023.105558

Shi, X.Q., Guo, W.C., Wan, P.J., Zhou, L.T., Ren, X.L., Ahmat, T., Fu, K.Y., Li, G.Q., 2013. Validation of reference genes for expression analysis by quantitative real-time PCR in *Leptinotarsa decemlineata* (Say). BMC Res Notes 6. 10.1186/1756-0500-6-93

Tebbe, C., Breckheimer, B., Racca, P., Schorn, C., Kleinhenz, B., Nauen, R., 2016. Incidence and spread of knockdown resistance kdr in German Colorado potato beetle (*Leptinotarsa decemlineata* Say) populations. EPPO Bulletin 46, 129–138. 10.1111/epp.12265

Van Leeuwen, T., Demaeght, P., Osborne, E.J., Dermauw, W., Gohlke, S., Nauen, R., Grbić, M., Tirry, L., Merzendorfer, H., Clark, R.M., 2012. Population bulk segregant mapping uncovers resistance mutations and the mode of action of a chitin synthesis inhibitor in arthropods. Proc Natl Acad Sci U S A 109, 4407–4412. 10.1073/pnas.1200068109

Wan, P.J., Shi, X.Q., Kong, Y., Zhou, L.T., Guo, W.C., Ahmat, T., Li, G.Q., 2013. Identification of cytochrome P450 monooxygenase genes and their expression profiles in cyhalothrin-treated Colorado potato beetle, *Leptinotarsa decemlineata*. Pestic Biochem Physiol 107, 360–368. 10.1016/j.pestbp.2013.10.004

Wickham, H., 2016. ggplot2. Springer International Publishing, Cham. 10.1007/978-3-319-24277-4

Wierenga, J.M., Hollingworth, R.M., 1994. The role of metabolic enzymes in insecticide-resistant Colorado potato beetles. Pestic Sci 40, 259–264. 10.1002/ps.2780400403

Wilhelm, L., Wang, Y., Xu, S., 2024. The Colorado potato beetle gene expression atlas. 10.1101/2024.03.28.587222

Williamson, M.S., Martinez-Torres, D., Hick, C.A., Devonshire, A.L., 1996. Identification of mutations in the housefly para-type sodium channel gene associated with knockdown resistance (kdr) to pyrethroid insecticides. Mol Gen Genet 252, 51–60. 10.1007/BF02173204

Xu, H., Pan, Y., Li, J., Yang, F., Chen, X., Gao, X., Wen, S., Shang, Q., 2022. Chemosensory proteins confer adaptation to the ryanoid anthranilic diamide insecticide cyantraniliprole in *Aphis gossypii* glover. Pestic Biochem Physiol 184. 10.1016/j.pestbp.2022.105076

Yan, J., Zhang, C., Zhang, M., Zhou, H., Zuo, Z., Ding, X., Zhang, R., Li, F., Gao, Y., 2023. Chromosome-level genome assembly of the Colorado potato beetle, *Leptinotarsa decemlineata*. Sci Data 10. 10.1038/s41597-023-01950-5

Yan, Y., Aumann, R.A., Häcker, I., Schetelig, M.F., 2023. CRISPR-based genetic control strategies for insect pests. J Integr Agric 22, 651–668. 10.1016/j.jia.2022.11.003

Ye, J., Coulouris, G., Zaretskaya, I., Cutcutache, I., Rozen, S., Madden, T.L., 2012. Primer-BLAST: A tool to design target-specific primers for polymerase chain reaction. BMC Bioinformatics 13, 134. 10.1186/1471-2105-13-134

Yu, S.J., Cong, L., Pan, Q., Ding, L.L., Lei, S., Cheng, L.Y., Fang, Y.H., Wei, Z.T., Liu, H.Q., Ran, C., 2021. Whole genome sequencing and bulked segregant analysis suggest a new mechanism of amitraz resistance in the citrus red mite, *Panonychus citri* (Acari: Tetranychidae). Pest Manag Sci 77, 5032–5048. 10.1002/ps.6544

Zhu, F., Moural, T.W., Nelson, D.R., Palli, S.R., 2016. A specialist herbivore pest adaptation to xenobiotics through up-regulation of multiple Cytochrome P450s. Sci Rep 6. 10.1038/srep20421

