## Supplemental for "Bulk segregant analysis reveals genomic regions associated with imidacloprid resistance in the Colorado potato beetle"

**Supplementary Information**

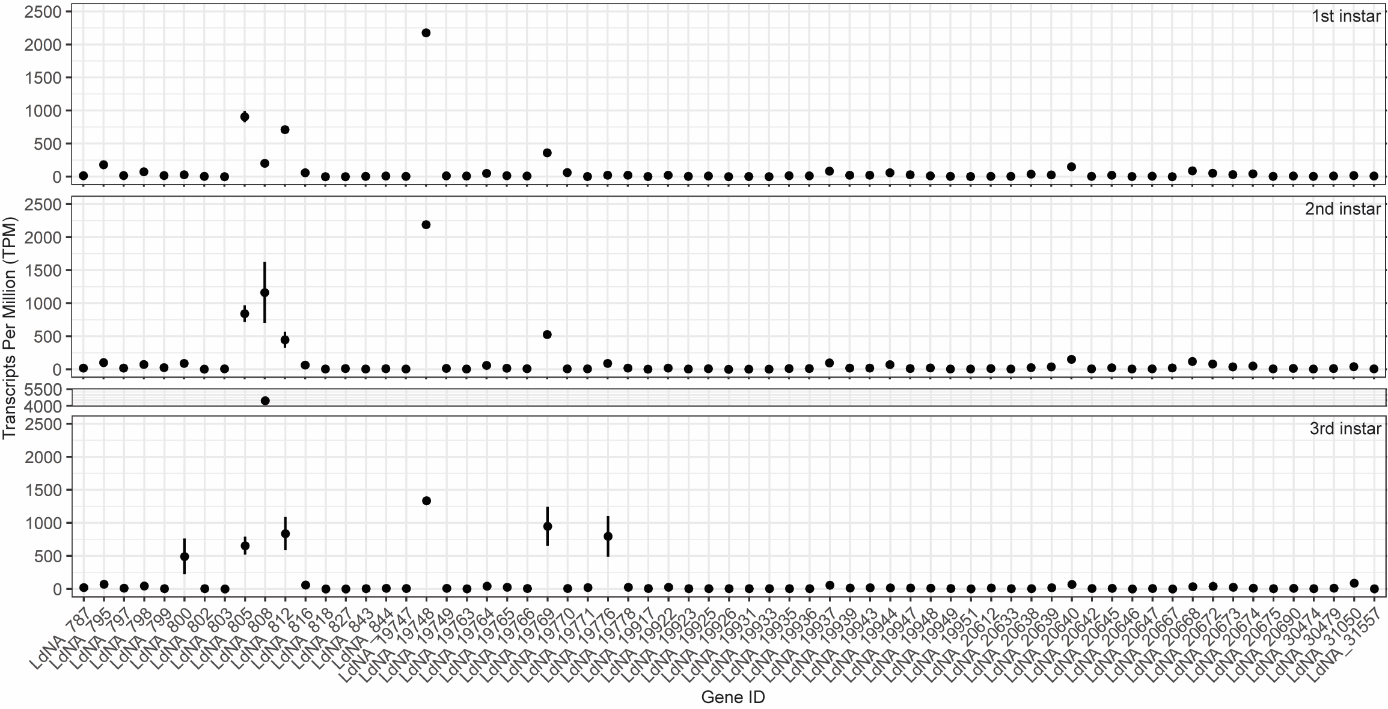

**Supplementary Figure 1. Expression profiles of 65 candidate genes from the CPB Gene Expression Atlas.** The graph shows the expression profiles of 65 candidate genes in Transcripts Per Million (TPM). The data was downloaded from the CPB GEA website (Wilhelm et al., 2024). The y-axis indicates the genes in the order of chromosome and position within the chromosome. The panels correspond to the first, second and third instar larvae as indicated on the top. Error bars show standard deviation.

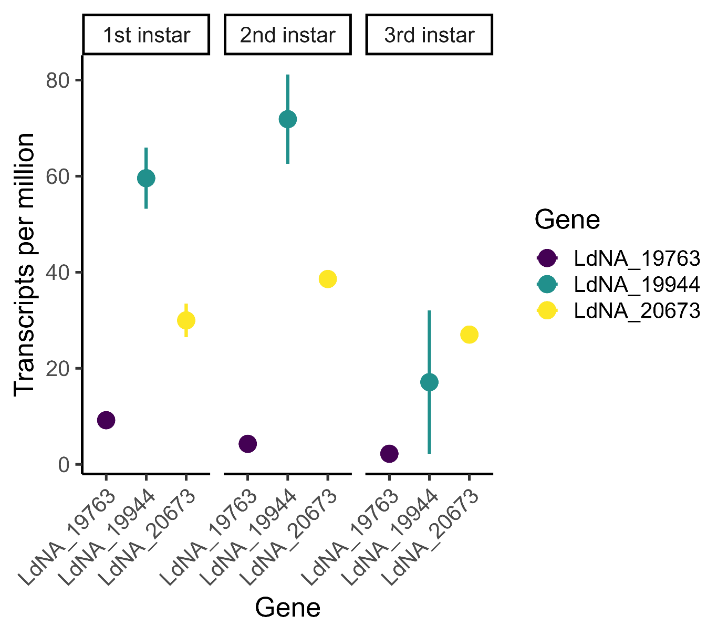
**Supplementary Figure 2. Expression profiles of three candidates from the CPB Gene Expression Atlas.** The graph shows the expression profiles of three candidate genes (LdNA19763, LdNA19944 and LdNA20673) in Transcripts Per Million (TPM). The data was downloaded from the CPB GEA website (Wilhelm et al., 2024). The colors indicate the different genes as indicated in the legend and the larval stage is shown on the top. Error bars show standard deviation.

**Supplementary Table 1. Primers used for RT-qPCR.**

| Gene | Forward (5’-3’) | Reverse (5’-3’) | Efficiency |
| --- | --- | --- | --- |
| RP18* | TAGAATCCTCAAAGCAGGTGGCGA | AGCTGGACCAAAGTGTTTCACTGC | 2.01 |
| ARF1* | CGGTGCTGGTAAAACGACAA | TGACCTCCCAAATCCCAAAC | 2.00 |
| LdNA19763 | AGAAGGGTCTTGTTGCGGC | GGCCTCGTGGGATATGAGAAG | 2.05 |
| LdNA19944 | AGTATCCAGGGGGCTACAGG | ATAAATGGCACGAAAGCCGC | 2.01 |
| LdNA20673 | GATGGGGAGGAGAAGATGATGA | AGTTGAGCCCATCATACCACTG | 2.08 |

*sequences obtained from Shi et al., 2013.

**Supplementary Table 2. dsRNA sequences used for RNAi**

| Gene | Sequence (5’-3’) | Size (bp) |
| --- | --- | --- |
| eGFP | 1 CACATGAAGC AGCACGACTT CTTCAAGTCC GCCATGCCCG AAGGCTACGT CCAGGAGCGC  61 ACCATCTTCT TCAAGGACGA CGGCAACTAC AAGACCCGCG CCGAGGTGAA GTTCGAGGGC  121 GACACCCTGG TGAACCGCAT CGAGCTGAAG GGCATCGACT TCAAGGAGGA CGGCAACATC  181 CTGGGGCACA AGCTGGAGTA CAACTACAAC AGCCACAACG TCTATATCAT GGCCGACAAG  241 CAGAAGAACG GCATCAAGGT GAACTTCAAG ATCCGCCACA ACATCGAGGA CGGCAGCGTG  301 CAGCTCGCCG ACCACTACCA GCAGAACACC CCCATCGGCG ACGGCCCCGT GCTGCTGCCC  361 GACAACCACT ACCTGAGCA | 379 |
| LdNA19763 | 1 CTAGTCTTCG GAGACATAGT GGGAGTTTTA GCTATTGAGG ACAACGAAAA GGTCAGATCA  61 GAAACGAATA TGTTTTGCCT GTATTTCCTG ATACTTGGAA TAGTTACTGG ACTAGCTACG  121 TTATTTCAGA TGTTCTCTTT CACTGTGGCA TGTGAGAAAC TGACATTGAG GCTGAGGAAG  181 AAGACATTCG AAGCCATGTT GAAACAAGAA ATGGGGTGGT TCGATAGGAA AGAAAATGGC  241 GTTGGAGCTT TGTGTGC | 257 |
| LdNA20673 | 1 GCAAAAATGA TGAATTATGG AGCTAAGGTG GCTATGAAAT TCCAGTATCA CTGCCTCATA  61 CTTCATGATG TAGATCTTAT TCCAATGAAT ACTGCAAATA TATATGGATG CACCAAATCG  121 CCTAGGCATA TGTCAAGTAG CTTGGATACA TTCAGATATA ACCTTCCATA CTTGACTCTC  181 TTCGGAGGAG CAGTTGCAAT TTCATCTGAA CAATTTCAGA AAGTCAATGG CATGTCCAAT  241 AGATTTTATG GATGGGGAGG AGAAGATG | 268 |

**Supplementary Table 3. The full list of genes identified in the significant peaks.** Genes identified in the genomic regions that are likely to be involved in imidacloprid resistance. The potential function of the gene is given in the ‘BLAST Annotation’ column if available; ‘-’ indicates lack of information. The ‘Start’ and ‘End’ columns indicate the position of the gene in bp within the specific chromosome.

| Sl. no. | Chr. | Start | End | Gene Name | Blast Annotation |
| --- | --- | --- | --- | --- | --- |
| 1 | 1 | 7475815 | 7479839 | LdNA_782 | Protein GPR107 |
| 2 | 1 | 7482998 | 7488169 | LdNA_783 | - |
| 3 | 1 | 7490021 | 7490615 | LdNA_784 | - |
| 4 | 1 | 7493715 | 7493984 | LdNA_785 | - |
| 5 | 1 | 7505195 | 7515539 | LdNA_786 | Probable G-protein coupled receptor B0563.6 |
| 6 | 1 | 7541214 | 7541843 | LdNA_787 | Slit homolog 3 protein |
| 7 | 1 | 7587150 | 7587800 | LdNA_788 | - |
| 8 | 1 | 7596910 | 7597363 | LdNA_789 | Large ribosomal subunit protein uL18 |
| 9 | 1 | 7605020 | 7606894 | LdNA_790 | PiggyBac transposable element-derived protein 3 |
| 10 | 1 | 7619210 | 7619530 | LdNA_791 | - |
| 11 | 1 | 7747602 | 7751589 | LdNA_792 | - |
| 12 | 1 | 7773326 | 7773960 | LdNA_793 | - |
| 13 | 1 | 7819634 | 7820104 | LdNA_794 | - |
| 14 | 1 | 7829263 | 7852833 | LdNA_795 | Zinc carboxypeptidase A 1 |
| 15 | 1 | 7837894 | 7842198 | LdNA_796 | Transposon Tf2-9 polyprotein |
| 16 | 1 | 7859085 | 7876267 | LdNA_797 | NEDD8-activating enzyme E1 regulatory subunit |
| 17 | 1 | 7869091 | 7870339 | LdNA_798 | Enoyl-CoA delta isomerase 1, mitochondrial |
| 18 | 1 | 7876684 | 7879733 | LdNA_799 | Bifunctional purine biosynthesis protein ATIC |
| 19 | 1 | 7879380 | 7894610 | LdNA_800 | Prostaglandin F synthase 1 |
| 20 | 1 | 7900576 | 7901418 | LdNA_801 | - |
| 21 | 1 | 7910994 | 7911822 | LdNA_802 | - |
| 22 | 1 | 7955891 | 7976740 | LdNA_803 | - |
| 23 | 1 | 7982065 | 7993943 | LdNA_804 | - |
| 24 | 1 | 8003077 | 8005511 | LdNA_805 | Cathepsin L-like proteinase |
| 25 | 1 | 8021203 | 8022263 | LdNA_806 | Transposable element P transposase |
| 26 | 1 | 8022438 | 8023463 | LdNA_807 | - |
| 27 | 1 | 8041483 | 8045590 | LdNA_808 | Cathepsin L-like proteinase |
| 28 | 1 | 8077520 | 8079996 | LdNA_809 | - |
| 29 | 1 | 8088672 | 8090807 | LdNA_810 | Cathepsin L-like proteinase |
| 30 | 1 | 8097070 | 8103597 | LdNA_811 | - |
| 31 | 1 | 8112169 | 8116310 | LdNA_812 | Crustapain |
| 32 | 1 | 8128862 | 8129777 | LdNA_813 | PiggyBac transposable element-derived protein 4 |
| 33 | 1 | 8130123 | 8131718 | LdNA_814 | - |
| 34 | 1 | 8145134 | 8151111 | LdNA_815 | Periaxin |
| 35 | 1 | 8221306 | 8274067 | LdNA_816 | Probable chitinase 10 |
| 36 | 1 | 8297926 | 8308745 | LdNA_817 | - |
| 37 | 1 | 8333172 | 8352457 | LdNA_818 | - |
| 38 | 1 | 8381028 | 8381975 | LdNA_819 | - |
| 39 | 1 | 8386622 | 8387596 | LdNA_820 | - |
| 40 | 1 | 8388202 | 8390009 | LdNA_821 | - |
| 41 | 1 | 8403624 | 8426005 | LdNA_822 | - |
| 42 | 1 | 8426640 | 8426912 | LdNA_823 | - |
| 43 | 1 | 8430604 | 8442624 | LdNA_824 | - |
| 44 | 1 | 8461513 | 8472674 | LdNA_825 | - |
| 45 | 1 | 8480053 | 8487581 | LdNA_826 | Galactokinase |
| 46 | 1 | 8491027 | 8499576 | LdNA_827 | - |
| 47 | 1 | 8552687 | 8553667 | LdNA_828 | - |
| 48 | 1 | 8558117 | 8565705 | LdNA_829 | - |
| 49 | 1 | 8580756 | 8581781 | LdNA_830 | - |
| 50 | 1 | 8588261 | 8588788 | LdNA_831 | Dynein axonemal heavy chain 8 |
| 51 | 1 | 8589165 | 8589584 | LdNA_832 | - |
| 52 | 1 | 8592961 | 8593974 | LdNA_833 | - |
| 53 | 1 | 8603306 | 8606187 | LdNA_834 | - |
| 54 | 1 | 8621258 | 8624575 | LdNA_835 | Plasma membrane calcium-transporting ATPase 3 |
| 55 | 1 | 8627982 | 8641068 | LdNA_836 | Zinc finger protein 347 |
| 56 | 1 | 8651251 | 8653067 | LdNA_837 | Zinc finger protein 57 |
| 57 | 1 | 8655000 | 8656291 | LdNA_838 | Zinc finger protein 782 |
| 58 | 1 | 8666512 | 8669165 | LdNA_839 | Zinc finger protein 26 |
| 59 | 1 | 8679198 | 8680736 | LdNA_840 | Zinc finger protein 782 |
| 60 | 1 | 8687026 | 8712605 | LdNA_841 | Zinc finger protein 782 |
| 61 | 1 | 8716416 | 8721772 | LdNA_842 | - |
| 62 | 1 | 8722199 | 8730280 | LdNA_843 | Gastrula zinc finger protein XlCGF57.1 (Fragment) |
| 63 | 1 | 8734421 | 8750822 | LdNA_844 | - |
| 64 | 1 | 8755056 | 8755406 | LdNA_845 | - |
| 65 | 1 | 8755947 | 8757092 | LdNA_846 | PiggyBac transposable element-derived protein 3 |
| 66 | 1 | 8757317 | 8759333 | LdNA_847 | Aldose reductase |
| 67 | 8 | 43242277 | 43256459 | LdNA_17297 | - |
| 68 | 8 | 43274063 | 43275326 | LdNA_17298 | - |
| 69 | 8 | 43332759 | 43333040 | LdNA_17299 | - |
| 70 | 8 | 43383672 | 43384085 | LdNA_17300 | Chimeric ERCC6-PGBD3 protein |
| 71 | 8 | 43384111 | 43384566 | LdNA_17301 | PiggyBac transposable element-derived protein 2 |
| 72 | 8 | 43384751 | 43385037 | LdNA_17302 | - |
| 73 | 8 | 43419512 | 43420434 | LdNA_17303 | - |
| 74 | 8 | 43473432 | 43473683 | LdNA_17304 | - |
| 75 | 8 | 43473702 | 43473968 | LdNA_17305 | - |
| 76 | 8 | 43481305 | 43483315 | LdNA_17306 | - |
| 77 | 8 | 43503655 | 43504674 | LdNA_17307 | - |
| 78 | 8 | 43505246 | 43505608 | LdNA_17308 | - |
| 79 | 8 | 43524078 | 43535567 | LdNA_17309 | - |
| 80 | 8 | 43553918 | 43554385 | LdNA_17310 | - |
| 81 | 8 | 43554615 | 43554959 | LdNA_17311 | Retrovirus-related Pol polyprotein from transposon TNT 1-94 |
| 82 | 8 | 43555335 | 43555610 | LdNA_17312 | - |
| 83 | 8 | 43613957 | 43616011 | LdNA_17313 | Pro-Pol polyprotein |
| 84 | 8 | 43618625 | 43619017 | LdNA_17314 | - |
| 85 | 8 | 43683625 | 43684179 | LdNA_17315 | KRAB-A domain-containing protein 2 |
| 86 | 8 | 43725014 | 43725439 | LdNA_17316 | - |
| 87 | 8 | 43749021 | 43749287 | LdNA_17317 | - |
| 88 | 10 | 1515829 | 1523803 | LdNA_19740 | Zinc finger protein 90 |
| 89 | 10 | 1522993 | 1523583 | LdNA_19741 | - |
| 90 | 10 | 1523620 | 1524045 | LdNA_19742 | - |
| 91 | 10 | 1536527 | 1536829 | LdNA_19743 | - |
| 92 | 10 | 1557324 | 1557650 | LdNA_19744 | - |
| 93 | 10 | 1609788 | 1610000 | LdNA_19745 | - |
| 94 | 10 | 1654775 | 1661472 | LdNA_19746 | - |
| 95 | 10 | 1698756 | 1700628 | LdNA_19747 | Syntaxin-6 |
| 96 | 10 | 1713214 | 1719534 | LdNA_19748 | Small ribosomal subunit protein uS7 |
| 97 | 10 | 1725625 | 1732713 | LdNA_19749 | - |
| 98 | 10 | 1754621 | 1757545 | LdNA_19750 | Zinc finger MYM-type protein 1 |
| 99 | 10 | 1795128 | 1795409 | LdNA_19751 | - |
| 100 | 10 | 1845348 | 1845733 | LdNA_19752 | - |
| 101 | 10 | 1904985 | 1905191 | LdNA_19753 | - |
| 102 | 10 | 1905807 | 1906213 | LdNA_19754 | - |
| 103 | 10 | 1906448 | 1906675 | LdNA_19755 | - |
| 104 | 10 | 1907002 | 1907247 | LdNA_19756 | - |
| 105 | 10 | 1976349 | 2106600 | LdNA_19757 | Hepatic leukemia factor |
| 106 | 10 | 2006577 | 2007053 | LdNA_19758 | - |
| 107 | 10 | 2037209 | 2037484 | LdNA_19759 | - |
| 108 | 10 | 2037631 | 2038060 | LdNA_19760 | - |
| 109 | 10 | 2064696 | 2064905 | LdNA_19761 | - |
| 110 | 10 | 2204669 | 2247524 | LdNA_19762 | ATP-dependent translocase ABCB1 |
| 111 | 10 | 2254329 | 2313006 | LdNA_19763 | ATP-dependent translocase ABCB1 |
| 112 | 10 | 2323030 | 2343199 | LdNA_19764 | Ubiquitin carboxyl-terminal hydrolase 5 |
| 113 | 10 | 2349139 | 2360544 | LdNA_19765 | Lysophosphatidylserine lipase ABHD12 |
| 114 | 10 | 2364307 | 2370090 | LdNA_19766 | - |
| 115 | 10 | 2377982 | 2382520 | LdNA_19767 | - |
| 116 | 10 | 2397331 | 2397588 | LdNA_19768 | - |
| 117 | 10 | 2409450 | 2423753 | LdNA_19769 | Inositol oxygenase |
| 118 | 10 | 2439301 | 2503688 | LdNA_19770 | Putative helicase mov-10-B.1 |
| 119 | 10 | 2571028 | 2571447 | LdNA_19771 | Lysozyme C-3 |
| 120 | 10 | 2592877 | 2593372 | LdNA_19772 | - |
| 121 | 10 | 2595959 | 2612625 | LdNA_19773 | - |
| 122 | 10 | 2648159 | 2648761 | LdNA_19774 | Jerky protein homolog-like |
| 123 | 10 | 2664127 | 2664783 | LdNA_19775 | PiggyBac transposable element-derived protein 2 |
| 124 | 10 | 2682578 | 2684445 | LdNA_19776 | Probable transaldolase |
| 125 | 10 | 2725183 | 2798848 | LdNA_19777 | Sex peptide receptor |
| 126 | 10 | 2803033 | 2920459 | LdNA_19778 | Rho guanine nucleotide exchange factor 12 |
| 127 | 10 | 7485132 | 7515401 | LdNA_19917 | Lactosylceramide 1,3-N-acetyl-beta-D-glucosaminyltransferase |
| 128 | 10 | 7525300 | 7525737 | LdNA_19919 | - |
| 129 | 10 | 7525300 | 7528778 | LdNA_19918 | - |
| 130 | 10 | 7587529 | 7598064 | LdNA_19920 | Band 7 protein AGAP004871 |
| 131 | 10 | 7600093 | 7600464 | LdNA_19921 | - |
| 132 | 10 | 7683600 | 7817876 | LdNA_19922 | G-protein coupled receptor Mth2 |
| 133 | 10 | 7734254 | 7734793 | LdNA_19923 | - |
| 134 | 10 | 7755935 | 7767274 | LdNA_19924 | - |
| 135 | 10 | 7779199 | 7779717 | LdNA_19925 | - |
| 136 | 10 | 7792849 | 7801228 | LdNA_19926 | - |
| 137 | 10 | 7829674 | 7830609 | LdNA_19927 | - |
| 138 | 10 | 7833588 | 7835213 | LdNA_19928 | Transposon Ty3-G Gag-Pol polyprotein |
| 139 | 10 | 7835335 | 7835610 | LdNA_19929 | - |
| 140 | 10 | 7838075 | 7838647 | LdNA_19930 | - |
| 141 | 10 | 7847991 | 7848506 | LdNA_19931 | - |
| 142 | 10 | 7855352 | 7856009 | LdNA_19932 | - |
| 143 | 10 | 7865598 | 7895072 | LdNA_19933 | G-protein coupled receptor Mth2 |
| 144 | 10 | 7898175 | 7898474 | LdNA_19934 | Tigger transposable element-derived protein 6 |
| 145 | 10 | 7899320 | 7916084 | LdNA_19935 | Probable ATP-dependent RNA helicase DDX52 |
| 146 | 10 | 7917349 | 7918317 | LdNA_19936 | Snurportin-1 |
| 147 | 10 | 7922417 | 7931189 | LdNA_19937 | Aspartate--tRNA ligase, cytoplasmic |
| 148 | 10 | 7931313 | 7931909 | LdNA_19938 | - |
| 149 | 10 | 7934588 | 7944525 | LdNA_19939 | Protein IWS1 homolog |
| 150 | 10 | 7936220 | 7939249 | LdNA_19940 | - |
| 151 | 10 | 7957395 | 7967542 | LdNA_19941 | - |
| 152 | 10 | 7999811 | 8012642 | LdNA_19942 | - |
| 153 | 10 | 8033082 | 8034442 | LdNA_19943 | UDP-glucuronic acid decarboxylase 1 |
| 154 | 10 | 8130421 | 8200857 | LdNA_19944 | ATP-dependent translocase ABCB1 |
| 155 | 10 | 8208802 | 8209710 | LdNA_19945 | - |
| 156 | 10 | 8212836 | 8214883 | LdNA_19946 | INO80 complex subunit E |
| 157 | 10 | 8220396 | 8221244 | LdNA_19947 | Probable E3 ubiquitin-protein ligase sinah |
| 158 | 10 | 8237205 | 8241177 | LdNA_19948 | - |
| 159 | 10 | 8261655 | 8311566 | LdNA_19949 | Probable G-protein coupled receptor Mth-like 3 |
| 160 | 10 | 8297839 | 8298156 | LdNA_19950 | - |
| 161 | 10 | 8320277 | 8359394 | LdNA_19951 | - |
| 162 | 10 | 8345085 | 8346473 | LdNA_19952 | Activity-regulated cytoskeleton associated protein 2 |
| 163 | 10 | 8365563 | 8367224 | LdNA_19953 | Gag-Pol polyprotein |
| 164 | 10 | 8369957 | 8370325 | LdNA_19954 | - |
| 165 | 10 | 8372440 | 8374541 | LdNA_19955 | Putative nuclease HARBI1 |
| 166 | 10 | 8374752 | 8382901 | LdNA_19956 | - |
| 167 | 10 | 8403620 | 8411768 | LdNA_19957 | - |
| 168 | 10 | 8447701 | 8460525 | LdNA_19958 | - |
| 169 | 10 | 29827485 | 29827805 | LdNA_20599 | - |
| 170 | 10 | 29828469 | 29829176 | LdNA_20600 | - |
| 171 | 10 | 29888885 | 29889208 | LdNA_20601 | - |
| 172 | 10 | 29911898 | 29912419 | LdNA_20602 | - |
| 173 | 10 | 29953444 | 29953662 | LdNA_20603 | - |
| 174 | 10 | 30071549 | 30073099 | LdNA_20604 | - |
| 175 | 10 | 30089856 | 30090224 | LdNA_20605 | - |
| 176 | 10 | 30114648 | 30116089 | LdNA_20606 | Chromatin assembly factor 1 p55 subunit |
| 177 | 10 | 30215549 | 30234445 | LdNA_20607 | - |
| 178 | 10 | 30248773 | 30252192 | LdNA_20608 | Facilitated trehalose transporter Tret1-1 |
| 179 | 10 | 30252230 | 30270727 | LdNA_20609 | - |
| 180 | 10 | 30268784 | 30269443 | LdNA_20610 | - |
| 181 | 10 | 30281483 | 30284806 | LdNA_20611 | - |
| 182 | 10 | 30286861 | 30288083 | LdNA_20612 | Facilitated trehalose transporter Tret1-2 homolog |
| 183 | 10 | 30357014 | 30358594 | LdNA_20613 | - |
| 184 | 10 | 30363727 | 30369833 | LdNA_20614 | - |
| 185 | 10 | 30390480 | 30391001 | LdNA_20615 | - |
| 186 | 10 | 30418329 | 30424575 | LdNA_20616 | Brother of CDO |
| 187 | 10 | 30443375 | 30443752 | LdNA_20617 | - |
| 188 | 10 | 30506791 | 30507084 | LdNA_20618 | - |
| 189 | 10 | 30509741 | 30516473 | LdNA_20619 | - |
| 190 | 10 | 30621651 | 30634135 | LdNA_20620 | - |
| 191 | 10 | 30670645 | 30671022 | LdNA_20621 | - |
| 192 | 10 | 30709741 | 30710115 | LdNA_20622 | - |
| 193 | 10 | 30775325 | 30779540 | LdNA_20623 | - |
| 194 | 10 | 30826596 | 30827087 | LdNA_20624 | - |
| 195 | 10 | 30870063 | 31077385 | LdNA_20625 | Alpha-catulin |
| 196 | 10 | 30964599 | 30964814 | LdNA_20626 | - |
| 197 | 10 | 31026846 | 31042062 | LdNA_20627 | Venom allergen 3 homolog |
| 198 | 10 | 31049621 | 31050142 | LdNA_20628 | 52 kDa repressor of the inhibitor of the protein kinase |
| 199 | 10 | 31050250 | 31050798 | LdNA_20629 | - |
| 200 | 10 | 31056225 | 31070536 | LdNA_20630 | - |
| 201 | 10 | 31086778 | 31090548 | LdNA_20631 | Serine/threonine-protein kinase Nek2 |
| 202 | 10 | 31093052 | 31115991 | LdNA_20632 | WD repeat-containing protein 3 |
| 203 | 10 | 31122297 | 31268581 | LdNA_20633 | Protein spitz |
| 204 | 10 | 31234068 | 31255743 | LdNA_20634 | - |
| 205 | 10 | 31439574 | 31439846 | LdNA_20635 | - |
| 206 | 10 | 31443982 | 31452867 | LdNA_20636 | - |
| 207 | 10 | 31473166 | 31473516 | LdNA_20637 | - |
| 208 | 10 | 31496992 | 31503626 | LdNA_20638 | Probable ribosome production factor 1 |
| 209 | 10 | 31510400 | 31526539 | LdNA_20639 | Retinoblastoma-binding protein 5 homolog |
| 210 | 10 | 31526591 | 31538840 | LdNA_20640 | Mitochondrial import receptor subunit TOM40 homolog 1 |
| 211 | 10 | 31548468 | 31548881 | LdNA_20641 | - |
| 212 | 10 | 31559226 | 31710217 | LdNA_20642 | Ras-GEF domain-containing family member 1B-A |
| 213 | 10 | 31733655 | 31734914 | LdNA_20643 | - |
| 214 | 10 | 31949502 | 31951571 | LdNA_20644 | Zinc finger protein 862 |
| 215 | 10 | 31953391 | 31954257 | LdNA_20645 | Distal membrane-arm assembly complex protein 2 |
| 216 | 10 | 31958730 | 32100639 | LdNA_20646 | Maternal embryonic leucine zipper kinase |
| 217 | 10 | 31975273 | 31983719 | LdNA_20647 | Cysteine-rich hydrophobic domain-containing protein 2 |
| 218 | 10 | 32023378 | 32024442 | LdNA_20648 | - |
| 219 | 10 | 32027839 | 32028456 | LdNA_20649 | G2/mitotic-specific cyclin-B |
| 220 | 10 | 32029894 | 32031024 | LdNA_20650 | G2/mitotic-specific cyclin-B |
| 221 | 10 | 32031640 | 32036151 | LdNA_20651 | G2/mitotic-specific cyclin-B |
| 222 | 10 | 32037260 | 32044470 | LdNA_20652 | G2/mitotic-specific cyclin-B |
| 223 | 10 | 32042402 | 32042788 | LdNA_20653 | - |
| 224 | 10 | 32161949 | 32162674 | LdNA_20654 | - |
| 225 | 10 | 32277523 | 32277894 | LdNA_20655 | - |
| 226 | 10 | 32287811 | 32288146 | LdNA_20656 | - |
| 227 | 10 | 32299092 | 32299643 | LdNA_20657 | - |
| 228 | 10 | 32301866 | 32303101 | LdNA_20658 | Pro-Pol polyprotein |
| 229 | 10 | 32309753 | 32310196 | LdNA_20659 | - |
| 230 | 10 | 32416126 | 32416590 | LdNA_20660 | Transposon Tf2-8 polyprotein |
| 231 | 10 | 32483179 | 32483427 | LdNA_20661 | - |
| 232 | 10 | 32559549 | 32561180 | LdNA_20662 | - |
| 233 | 10 | 32601497 | 32611590 | LdNA_20663 | - |
| 234 | 10 | 32660334 | 32660597 | LdNA_20664 | Large ribosomal subunit protein P1 |
| 235 | 10 | 32665516 | 32666388 | LdNA_20665 | KRAB-A domain-containing protein 2 |
| 236 | 10 | 32729211 | 32844399 | LdNA_20666 | Paired box protein Pax-2-A |
| 237 | 10 | 32877990 | 32882425 | LdNA_20667 | - |
| 238 | 10 | 32893983 | 32932389 | LdNA_20668 | - |
| 239 | 10 | 32948110 | 32950568 | LdNA_20669 | - |
| 240 | 10 | 32973204 | 32991908 | LdNA_20670 | Chaoptin |
| 241 | 10 | 33070580 | 33072086 | LdNA_20671 | - |
| 242 | 10 | 33146685 | 33180110 | LdNA_20672 | Protein transport protein Sec24C |
| 243 | 10 | 33182649 | 33194725 | LdNA_20673 | Beta-1,4-galactosyltransferase 1 |
| 244 | 10 | 33196991 | 33203422 | LdNA_20674 | - |
| 245 | 10 | 33228756 | 33229043 | LdNA_20675 | - |
| 246 | 10 | 33349915 | 33350147 | LdNA_20676 | - |
| 247 | 10 | 33350174 | 33350383 | LdNA_20677 | - |
| 248 | 10 | 33358044 | 33358247 | LdNA_20678 | - |
| 249 | 10 | 33461716 | 33462960 | LdNA_20679 | Retrovirus-related Gag polyprotein from transposon HMS-Beagle |
| 250 | 10 | 33471162 | 33471476 | LdNA_20680 | - |
| 251 | 10 | 33515717 | 33516085 | LdNA_20681 | - |
| 252 | 10 | 33555774 | 33556073 | LdNA_20682 | Small ribosomal subunit protein eS12 |
| 253 | 10 | 33602851 | 33604584 | LdNA_20683 | - |
| 254 | 10 | 33648567 | 33650675 | LdNA_20684 | PiggyBac transposable element-derived protein 4 |
| 255 | 10 | 33715181 | 33715887 | LdNA_20685 | - |
| 256 | 10 | 33722458 | 33726962 | LdNA_20686 | - |
| 257 | 10 | 33731705 | 33732228 | LdNA_20687 | - |
| 258 | 10 | 33829300 | 33829932 | LdNA_20688 | - |
| 259 | 10 | 33830509 | 33831022 | LdNA_20689 | - |
| 260 | 10 | 34061079 | 34069751 | LdNA_20690 | - |
| 261 | 10 | 34187175 | 34190390 | LdNA_20691 | - |
| 262 | 10 | 34289362 | 34297239 | LdNA_20692 | - |
| 263 | 10 | 34304316 | 34337932 | LdNA_20693 | - |
| 264 | 10 | 34328299 | 34337932 | LdNA_20694 | Dynein regulatory complex protein 9 |
| 265 | 16 | 5560257 | 5560562 | LdNA_30459 | - |
| 266 | 16 | 5584075 | 5584626 | LdNA_30460 | - |
| 267 | 16 | 5588942 | 5646670 | LdNA_30461 | - |
| 268 | 16 | 5637284 | 5637865 | LdNA_30462 | - |
| 269 | 16 | 5638974 | 5639564 | LdNA_30463 | Transposon Tf2-9 polyprotein |
| 270 | 16 | 5640646 | 5641071 | LdNA_30464 | - |
| 271 | 16 | 5656599 | 5670615 | LdNA_30465 | Protein rhomboid |
| 272 | 16 | 5671820 | 5672083 | LdNA_30466 | - |
| 273 | 16 | 5672152 | 5672837 | LdNA_30467 | RNA-directed DNA polymerase from mobile element jockey |
| 274 | 16 | 5704985 | 5705533 | LdNA_30468 | Retrovirus-related Pol polyprotein from transposon 17.6 |
| 275 | 16 | 5706399 | 5706680 | LdNA_30469 | - |
| 276 | 16 | 5727953 | 5821831 | LdNA_30470 | REPTOR-binding partner |
| 277 | 16 | 5738012 | 5738257 | LdNA_30471 | - |
| 278 | 16 | 5829423 | 5829731 | LdNA_30472 | - |
| 279 | 16 | 5832969 | 5833385 | LdNA_30473 | - |
| 280 | 16 | 5836460 | 5968027 | LdNA_30474 | DENN domain-containing protein 2B |
| 281 | 16 | 5860803 | 5861222 | LdNA_30475 | HEAT repeat-containing protein 6 |
| 282 | 16 | 5862624 | 5863217 | LdNA_30476 | HEAT repeat-containing protein 6 |
| 283 | 16 | 5959689 | 5960072 | LdNA_30477 | - |
| 284 | 16 | 5979974 | 6042756 | LdNA_30478 | Vigilin |
| 285 | 16 | 6052497 | 6187195 | LdNA_30479 | Fibrillin-1 |
| 286 | 16 | 6185388 | 6185741 | LdNA_30480 | - |
| 287 | 16 | 22159527 | 22181417 | LdNA_31050 | Phenoloxidase 1 |
| 288 | 16 | 22191322 | 22191687 | LdNA_31051 | - |
| 289 | 16 | 22192538 | 22192849 | LdNA_31052 | - |
| 290 | 16 | 22203882 | 22267213 | LdNA_31053 | Leucine-rich repeat-containing protein 15 |
| 291 | 16 | 22238249 | 22238470 | LdNA_31054 | - |
| 292 | 16 | 31567397 | 31651247 | LdNA_31513 | Anoctamin-10 |
| 293 | 16 | 31592253 | 31593505 | LdNA_31514 | Probable E3 ubiquitin-protein ligase sinah |
| 294 | 16 | 31597561 | 31598595 | LdNA_31515 | Probable E3 ubiquitin-protein ligase sinah |
| 295 | 16 | 31602894 | 31604147 | LdNA_31516 | Probable E3 ubiquitin-protein ligase sinah |
| 296 | 16 | 31607476 | 31608722 | LdNA_31517 | Probable E3 ubiquitin-protein ligase sinah |
| 297 | 16 | 31630005 | 31635496 | LdNA_31518 | - |
| 298 | 16 | 31643517 | 31643930 | LdNA_31519 | Chimeric ERCC6-PGBD3 protein |
| 299 | 16 | 31696096 | 31768943 | LdNA_31520 | Periostin |
| 300 | 16 | 31770678 | 31773106 | LdNA_31521 | PITH domain-containing protein GA19395 |
| 301 | 16 | 31774061 | 31777250 | LdNA_31522 | Calmodulin-like protein 4 |
| 302 | 16 | 31781246 | 31789010 | LdNA_31523 | Mitochondrial coenzyme A transporter SLC25A42 |
| 303 | 16 | 31791663 | 31800811 | LdNA_31524 | Coiled-coil domain-containing protein 6 |
| 304 | 16 | 31805645 | 31808711 | LdNA_31525 | - |
| 305 | 16 | 31812353 | 31815338 | LdNA_31526 | - |
| 306 | 16 | 31819369 | 31848927 | LdNA_31527 | Stathmin |
| 307 | 16 | 31825792 | 31832355 | LdNA_31528 | - |
| 308 | 16 | 31851067 | 31854946 | LdNA_31529 | Probable serine hydrolase |
| 309 | 16 | 31857669 | 31865250 | LdNA_31530 | Probable serine hydrolase |
| 310 | 16 | 31872431 | 31876121 | LdNA_31531 | Probable serine hydrolase |
| 311 | 16 | 31876461 | 31878803 | LdNA_31532 | Solute carrier family 25 member 44 |
| 312 | 16 | 31884287 | 31903796 | LdNA_31533 | Signal transducer and activator of transcription 5B |
| 313 | 16 | 31904946 | 31910017 | LdNA_31534 | Zinc finger and SCAN domain-containing protein 22 |
| 314 | 16 | 31910855 | 31912536 | LdNA_31535 | Lysosomal Pro-X carboxypeptidase |
| 315 | 16 | 31914346 | 31914945 | LdNA_31536 | - |
| 316 | 16 | 31917619 | 31918218 | LdNA_31537 | - |
| 317 | 16 | 31919467 | 31920293 | LdNA_31538 | - |
| 318 | 16 | 31920863 | 31921462 | LdNA_31539 | - |
| 319 | 16 | 31994681 | 31995883 | LdNA_31540 | FERM domain-containing protein 5 |
| 320 | 16 | 32016722 | 32113481 | LdNA_31541 | Zinc finger protein Gfi-1 |
| 321 | 16 | 32129153 | 32129960 | LdNA_31542 | General odorant-binding protein 19d |
| 322 | 16 | 32131486 | 32131761 | LdNA_31543 | - |
| 323 | 16 | 32141306 | 32141959 | LdNA_31544 | PiggyBac transposable element-derived protein 3 |
| 324 | 16 | 32142340 | 32142549 | LdNA_31545 | - |
| 325 | 16 | 32157017 | 32157436 | LdNA_31546 | - |
| 326 | 16 | 32163337 | 32170971 | LdNA_31547 | Small ribosomal subunit protein eS10B |
| 327 | 16 | 32195099 | 32196341 | LdNA_31548 | Small ribosomal subunit protein uS5 |
| 328 | 16 | 32205509 | 32209034 | LdNA_31549 | - |
| 329 | 16 | 32217775 | 32220465 | LdNA_31550 | Lysosomal thioesterase PPT2 homolog |
| 330 | 16 | 32221334 | 32222854 | LdNA_31551 | Origin recognition complex subunit 2 |
| 331 | 16 | 32223102 | 32224220 | LdNA_31552 | Transcription initiation factor TFIID subunit 10 |
| 332 | 16 | 32231032 | 32238119 | LdNA_31553 | - |
| 333 | 16 | 32267087 | 32267782 | LdNA_31554 | Fumarylacetoacetate hydrolase domain-containing protein 2A |
| 334 | 16 | 32267882 | 32270368 | LdNA_31555 | Fumarylacetoacetate hydrolase domain-containing protein 2 |
| 335 | 16 | 32273633 | 32275078 | LdNA_31556 | - |
| 336 | 16 | 32275644 | 32278200 | LdNA_31557 | - |
| 337 | 16 | 32278225 | 32279055 | LdNA_31558 | WD repeat and FYVE domain-containing protein 3 |
